# On the existence of mechanoreceptors within the neurovascular unit of the rodent and rabbit brain

**DOI:** 10.1101/480921

**Authors:** Jorge Larriva-Sahd, Martha León-Olea, Víctor Vargas-Barroso, Alfredo Varela-Echavarría, Luis Concha

## Abstract

We describe a set of perivascular interneurons (PINs) originating a series of fibro-vesicular complexes (FVCs) throughout the gray matter of the adult rabbit and rat brain. PINs-FVCs are ubiquitous throughout the brain vasculature as defined in Golgi-impregnated specimens. Most PINs consist of small, aspiny cells with local or long (> 1 millimeter) axons that split running with arterial blood vessels. Upon ramification, axons originate FVCs around the roots of the arising vascular branches. Distally, FVCs form paired axons that run parallel to the vessel’s wall until another ramification ensues and a new FVC is formed. This alternating pattern ceases when the capillary diameter narrows (i.e., <8 µm) and axons resolve. FVCs, as visualized by electron microscopy, consist of clusters of anastomotic perivascular bulbs (PVBs) arising from the PIN’s unmyelinated axon. PVBs lie alongside the pre- or -capillary wall, surrounded by end-feet and the extracellular matrix of endothelial cells and pericytes. A PVB contains mitochondria, multivesicular bodies, and granules with a membranous core similar to those observed in Meissner corpuscles and other mechanoreceptors. Some PVBs form asymmetrical, axo-spinous synapses with presumptive adjacent neurons. Antisera to sensory fiber-terminals co-label putative FVCs that are embedded by astrocytic end-feet. Because of the strategic location, ubiquity, and cytological organization of the PIN-FVC, it is suggested that: 1. PIN-FVCs are distributed throughout the mammalian brain vasculature. 2. The PIN-FVC is a putative sensory receptor intrinsic of the neurovascular unit. 3. The PIN-FVC may correspond to an afferent limb of the sensory-motor feed-back controlling local blood flow.

## 1. Introduction

The mechanisms underpinning cerebral blood flow pertain to both basic and applied neuroscience. In this broad context, homeostatic blood supply to active areas of the brain is dependent on a complex neuro-vascular response. Two interrelated mechanisms provide a continuous and adjusting perfusion: a constant cerebral blood flow over a wide-range of arterial pressure and local functional hyperaemia (FH)(Idecola and Nedergaard, 2007, Filosa, et al. 2016). Arterial blood pressure results from the orchestrated action of the cardiac output, blood viscosity, and muscular tone of the arteries to maintain it within a physiological range. This dynamic process is monitored by mechanoreceptors-and chemo-receptors that are associated with large arteries and decode both blood pressure and composition. Central convergence of peripheral receptors onto ganglion cells and then onto cranial nerve nuclei results in adaptive vegetative motor responses (Hall, 2016). Distally, FH results from the coordinated action of arterioles and arterial capillaries with the neighboring glial and neural elements, structuring the *neurovascular unit* (NVU) (Roy and Sherrington 1890). While there is a general agreement about the synergistic involvement of the vascular, glial, and neural elements in detecting FH, it is unclear how mechanical or chemical information arising from the blood flow is transduced for an eventual vasomotor feed-back response (see Busse and Fleming 2003; Zonta, 2003; Idecola 2004; Filosa et al. 2016). Relevant to the present study is the distinct group of cortical interneurons reactive to nitric acid diaphorase (NADPH-p) and/or neuronal nitric oxide-synthase (nNOS-I) that has been identified (Estrada and DeFelipe 1998; Vaucher et al. 2000; Ducheim et al. 2012). Interestingly, axons of a group of cortical NADPH-p interneurons surround cortical blood vessels suggesting that these cells are implicated in local mechanisms controlling FH (Estrada and DeFelipe 1998; Ducheim et al. 2012). Furthermore, given that nNOS-I neurons in the myenteric plexus are selectively activated by colonic elongation, they have been termed *mechanoreceptor neurons* (see Smith et al. 2007). Due to the absence of cellular substrate(s) transducing mechanical cues to receptor potentials (RPs) from the blood stream to the neighboring neural parenchyma (Idecola 2004; Filosa et al. 2016) systematic search for cellular substrate(s) is required. The Golgi technique revealed a unique group of small perivascular interneurons (PINs) associated with arterial blood vessels throughout the brain parenchyma. A set of PINs issues long axons that proceed counter-current along the arterial blood vessel organizing into fibro-vesicular complexes (FVCs) that encircle the roots of tributary blood vessels. Combined electron microscopic observations and 3-D reconstructions at sites of blood vessel ramification revealed that the FVC is composed of perivascular bulbs (PVBs) encased by astrocytic end-feet (EF) and capillary basal laminae. This association, coupled with the membrane-bound organelles contained in the PVB resembling those of the Meissner corpuscle and other mechanoreceptors (Andres and During 1973; Chouchkov 1973; Hashimoto 1973; Ide et al. 1987), suggests that the PIN-FVCs is involved in decode mechanical cues arising from the brain’s arterial and capillary blood flow. Immunocytochemical work featuring putative neurotransmitters and 3D interactions between these presumptive receptors will be presented separately.

## 2. Materials and Methods

### 2.1 Animals

Adult rats, rabbits and mice raised in our pathogen-free vivarium were utilized. Animal manipulations and sacrifices obeyed ethical policies defined by our ad hoc Animal Research Committee.

### 2.2 Rapid Golgi Technique

Brains from normal adult albino rats (n=250) and New-Zealand rabbits (n=40) raised in our vivarium facility were studied. Following sacrifice with barbiturate overdose, brains were removed from the skull and transversely cut into four quarters with a pair of sharp scissors. From rabbit blocks of tissue approximating 4mm in thickness were sampled from the brain stem; cerebellum; frontal, parietal, and occipital isocortex; hippocampus-entorhinal cortex; diencephalon; basal ganglia; olfactory cortex; olfactory bulb; and peduncle. Aside from the species source, each block was placed for 10 to 13 days in an aqueous solution containing 0.25% osmium tetroxide and 3% potassium dichromate and transferred to a 0.75% silver nitrate solution for the following 20 days. Tissue blocks were held with a paraffin shell and sectioned at 150 microns with a sliding microtome. The sections were transferred to 70% ethyl-alcohol, dehydrated, cleared in Terpineol-xylene, mounted, and coverslipped with Entellan (Merck). Illustrations were obtained with a camera lucida adapted to an Axioplan 2 (Zeiss, Oberkochen) light microscope, using 40X and 100X oil objectives. Then, Indian ink and soft pencil replicas from both species provided demonstrative images. Somatic, axonal, dendritic, and vesicular structures were measured in digital images acquired with an AxioCam camera aided by Kontron software (Zeiss, Oberkochen).

### 2.3 Immunohistochemistry and Histochemistry

The primary antisera (Table 1) aimed to label sensory fibers included calcitonin gene-related peptide (CGRP) (Alvarez et al. 1993; Vega et al. 1996, Ishida-Yamamoto, et al. 1988, Johansson et al. 1999; Warfvinge and Edvisson, 2017) and vesicular glutamate co transporter 1 (VGlu1) (see Bewick, 2015; Varela-Echevarría et al. 2017) and were used as follows. Rat sections were left overnight in polyclonal antisera to CGRP (1:1000) and VGlu 1 (1:5000) dissolved in 0.1 M phosphate buffer (PB). Immunoreactive sites were revealed with the ABC kit according to the previously described protocol (Vargas-Barroso and Larriva-Sahd 2013). Supplemental sections from three mice of the colony *FVB/N-Tg (GFAP-EGFP)GFEA-FKi* that displayed EGFP (eGFP) fluorescence in astrocytes from multiple areas of the CNS (see Varela-Echevarría et al. 2017) were incubated with the antisera to CGRP (1:1000) and VGlu1 (1:500). Sections from both species were then sequentially immersed in a secondary rabbit Cy5 diluted 1:1000 in PB 0.8% saline (PB-S) for 2 h (Table 2). Following a 5-min wash in PBS, sections were incubated in DAPI diluted 1:4000 for 5 min and mounted in Mowiol medium. A set of adult rat sections was separately incubated with a primary antibody to nNOS. Following an overnight incubation with the primary antibody (Table 1), sections were processed with the protocol just described, visualizing immunoreactive sites with a donkey anti-rabbit antibody bound with Alexa fluor 555.

**Table 1.**
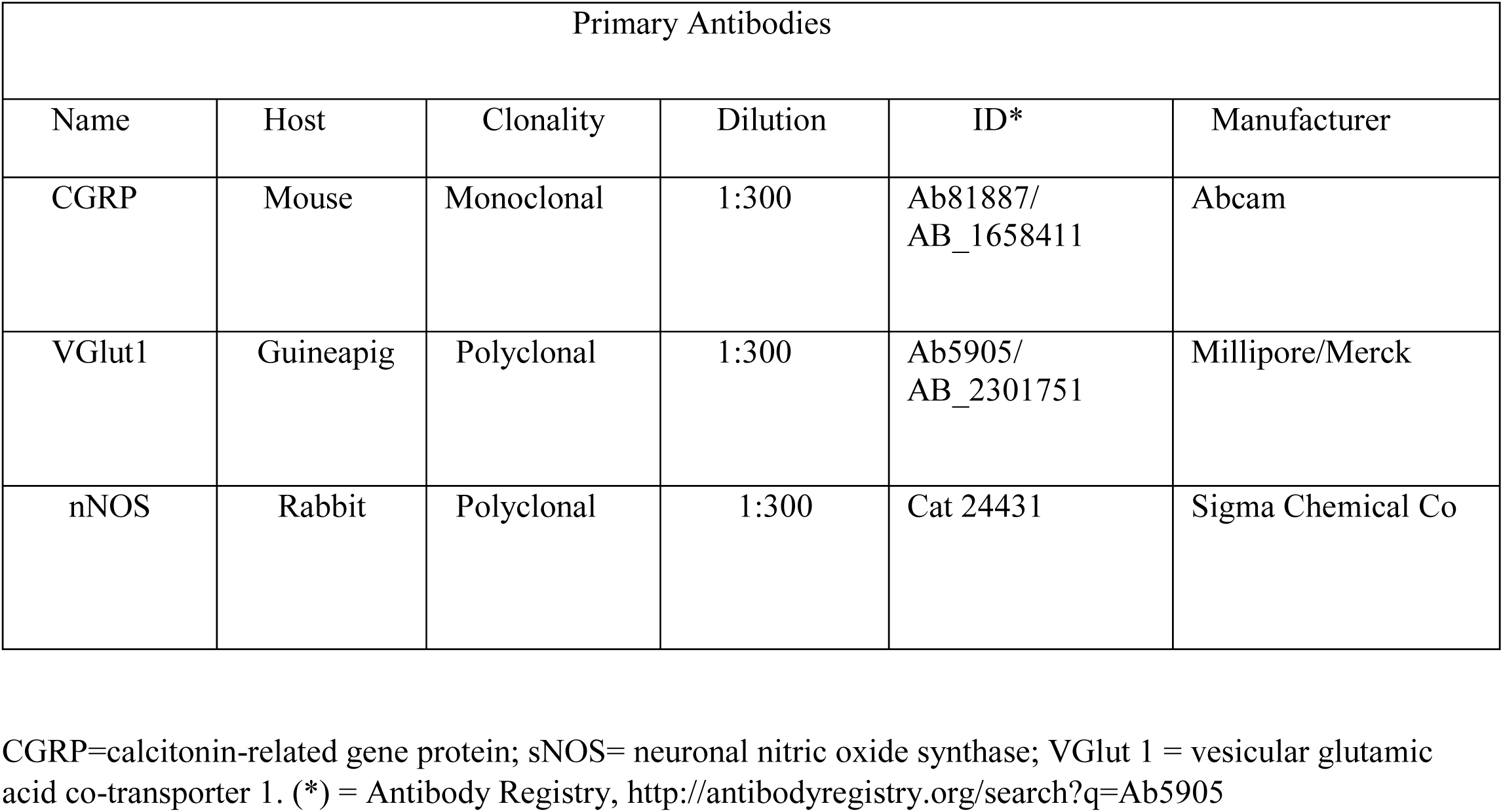
Antisera and reagents for immuno- and histo-fluorescence

| Primary Antibodies |  |  |  |  |  |
| --- | --- | --- | --- | --- | --- |
| Name | Host | Clonality | Dilution | ID* | Manufacturer |
| CGRP | Mouse | Monoclonal | 1:300 | Ab81887/<br>AB_1658411 | Abcam |
| VGlut1 | Guineapig | Polyclonal | 1:300 | Ab5905/<br>AB_2301751 | Millipore/Merck |
| nNOS | Rabbit | Polyclonal | 1:300 | Cat 24431 | Sigma Chemical Co |

CGRP=calcitonin-related gene protein; nNOS= neuronal nitric oxide synthase; VGlut 1 = vesicular glutamic acid co-transporter 1. (\*) = Antibody Registry, http://antibodyregistry.org/search?q=Ab5905

**Table 2.** Secondary antibodies

| Secondary Antibodies |  |  |  |  |  |  |
| --- | --- | --- | --- | --- | --- | --- |
| Name | Host | Fluorophore | EX/EM (nm) | Dilution | ID* | Manufacturer |
| Anti-Mouse | Goat | Alexa 350 | 345/440 | 1:1000 | A--- 11045<br>AB_142754 | Molecular Probes |
| Anti-Mouse | Donkey | Alexa Fluor 594 | 590/617 | 1:1000 | Ab150112 | Abcam |
| Anti-Guinea pig | Goat | Cy3 | 552/565 | 1:1000 | Ab102370<br>AB_10711466 | Abcam |
| Anti--Sheep | Donkey | Alexa Fluor594 | 590/617 | 1:1000 | Ab150180<br>AB_2716768 | Abcam |
| Anti-rabbit | Donkey | Alexa Fluor488 | 555 | 1:300 | A 31572 | Invitrogen Co |

(\*) = Antibody Registry) http://antibodyregistry.org/search?q=Ab5905

Three additional adult rats (n=3) and four GFAP-EGFP (eGFP) mice were anesthetized and perfused (see above) with 4% paraformaldehyde dissolved in 0.1M PB. Brains were sectioned with a vibratome at 40 microns and transferred to PB (see Varela-Echevarria et al. 2017). To visualize the B4 isolectin (IB4) (Table 3) that labels unmyelinated fibers (Silverman and Kruger 1990; see Eftekhari and Evinsson 2011) a set of sections were separately processed to visualize the NADPH-d reactivity. Floating sections were first immersed in PB solution containing 0.1% Triton X-100 for 5 min and then transferred to 0.1mg/ml nitro blue tetrazolium and 1mg of β-NADPH dissolved in PB for 45 min at room temperature. Following a rapid transfer to fresh PB, sections were mounted in glass slides, air-dried, and coversliped (Sánchez-Islas and León-Olea 2001).

**Table 3.** Reagents for Histochemistry

| Lectin |  |  |  |  |  |  |  |
| --- | --- | --- | --- | --- | --- | --- | --- |
| Name | Full Name | Species | Conjugate | EX/EM | Dilution | Product | Manufacturer |
| IB4 | Isolectin GS- IB4- Cy5 | *Griffonia*<br>*Simplicifolia* | Cy5 | 650/668 nm | 1:80 | 132450 | Invitrogen |

Observations and image acquisition were performed using a Zeiss 780 LSM confocal microscope. Confocal digital micrographs were acquired at a 0.30 µm interval with a 1,024 × 1,024 pixels size were further processed and edited with the Image J and Adobe Photoshop software, respectively. Illustrative 3D images were obtained with the Amira software.

### 2.4 Electron Microscopy

Ten male albino rats (n=5) were processed for routine electron microscopy as described elsewhere (Larriva-Sahd 2006). Each animal was sacrificed with an overdose of sodium pentobarbital (i.e., 30 mg/kg). Following perfusion through the left ventricle with 250ml of 2.5/4% glutaraldehyde/paraformaldehyde dissolved in 0.1M sodium cacodylate buffer, the brain was free-hand sectioned along the coronal plane and sampled according to the Swanson atlas (Swanson 2004) as follows. Parietal cortex: Level (L) 7. Horizontal (H) = 2.1 to 3.0 and vertical (V) = 0.8 to 2.0; frontal cortex: L8. H = 1.0 to 2.8, V = 1.0 to 3.0; piriform cortex: L2. H = 4.3 to 6, V = 9.6 to 10.2; anterior thalamus: L27. H = 1.0 to 3.9, V = 4.2 to 7; posterior thalamus: L33. H = 0.5 to 4.5, V = 4.5 to 8.0; and olfactory bulb medulla: L3. H = 0.8 to 1.2, V = 4 to 5.6. Tissue slices were postfixed for an hour in 1 % osmium tetra-oxide dissolved in the same buffer, dehydrated in acetone and flat-embedded in Epon. For light microscopy, one-micrometer-thin sections were obtained with a Leica microtome equipped with glass knives and stained with aqueous 0.5% toluidine blue. Vascular areas were identified at the light microscopic level, and then the block surface of the ultra-thin sections was reduced to include capillaries arising from larger (>80 µm in diameter) arteries for electron microscopic observations. Ultra-thin sections (60 to 70 nm) were mounted in single-slot copper grids that were previously covered with Formvar film. A JEOL 1010 electron microscope operated at 80 kV and equipped with a Gatan digital camera was used for observation. Following ultrastructural identification of FVCs associated with the capillary wall, the tissue blocks were serially sectioned with a Leica microtome with aided by a diamond knife. 3D reconstructions were performed with digital micrographs from a series of 50 to 190 sections that were first aligned with the Fuji software and then assembled with the Reconstruct software. Two to five reconstructions per brain area were obtained. To asses organelle homology between the vesicular structures described here and those from peripheral mechanoreceptors, additional palmar and digital skin samples from the forelimb were studied.

## 3. Results

### 3.1 Rapid Golgi Technique

The following is an account of the general features of PINs whose axons originate FVCs throughout the brain and brain stem gray matter of adult rats and rabbits. Due to the superior impregnation obtained from rabbit specimens, neuronal classification and axonal specializations are described in this species (Table 4); qualitative differences with respect to the rat specimens are highlighted.

**Table 4.** Golgi technique. Numbers, subtypes, and distribution of perivascular neurons in the adult rabbit

|  | sPIN | psPIN | bPIN | pPIN |
| --- | --- | --- | --- | --- |
| Isocortex | 5 | 5 | 4 | 4 |
| Hippocampus | 5 | 4 | 3 | 3 |
| Thalamus | 3 | 5 | 3 | 3 |
| Piriform cortex | 6 | 4 | 4 | 5 |
| Midbrain | 1 | 2 | 1 | 1 |
| Pons | 3 | 1 | 3 | 1 |
| Medulla | 2 | 1 | 3 | 1 |
| Cervical Spinal Cord | 1 | - | 3 | - |
| Total = 90 | 26 | 22 | 24 | 18 |

central nervous system.
PIN = perivascular neuron: b = bipolar; p = polygonal; ps = pseudomonopolar; s = spindle-shaped

The PIN’s soma commonly lies at sites of blood vessel ramification or, less commonly, along the shaft of the blood vessel throughout the central nervous system (CNS)(Table 4). An important characteristic of the PIN is that both the soma and its processes are distributed within the perivascular domain. The somata may be spindle-shaped (sPIN), bipolar (bPIN), pseudounipolar (psPIN), and polygonal (pPIN), in order of frequency (Table 4). The sPIN (Figs.1, 2a-e) bears a narrow perikaryon issuing a long axon and a rudimentary dendrite coursing in the opposite direction. After a short trajectory (<50 µm), the parent axon divides into paired or *twin* fibers (TFs), each one usually providing thin, recurrent branches. A striking characteristic of TFs is that upon ramification of the blood vessel, one or both fibrils sprout into an FVC that distributes within the parent and root of the arising blood vessel (Figs. 1a-c, 2a-e). Distal from the site of vascular forking, the FVC converges into twin fibrils that keep running until the blood vessel ramifies again and another FVC is formed. Implicitly, the number of TF-FVC intervals varies as a function of both the frequency of vascular forking and degree of parallelism between the blood vessel and the plane of sectioning. In privileged sections, TFs may be traced until they resolve into tuberosities underneath of the pial-parenchymal interphase, the so-called Virchow-Robin *space* (see Jones 1970). Hence, the full picture of the sPIN and its axon is visualized in the cerebral isocortex (Fig. 2a-e, h), hippocampus (Fig. 4a-d), olfactory bulb (not shown), and basal ganglia (Fig. 1), where long arteries follow a straight trajectory (Zhang et al. 2018). Elsewhere, in the cervical spinal cord (Online resource, Fig. 1), or brain stem (Fig. 2f, g, 4e, f) (Online Resource, Fig. 2), the blood vessels are shorter or course undulating paths; hence, only parts of the sPIN and pPIN are accessible to the observer (1). The bPIN soma also lays at the mesangial interstitium and it bears short processes that distribute therein (Fig. 2f). Commonly, the bPIN axon ramifies shortly afterwards (<40 µm) to give rise to FVC that surrounds the wall of one or two capillaries (see below). The bPIN’s single dendrite is short (<70µm), varicose, and with scarce spines. The third set of PINs, or psPINs (Fig 2g), have a rounded, egg-shaped soma originating a single short (<30 µm) process that divides into an axon and a dendrite. Both the psPIN and its processes gather those described for ganglion cells in the peripheral and CNS. The last type of neuron or pPIN is characterized by its relatively large perikaryon, and for providing 2-4, varicose dendrites and an axon that collateralizes first within the underlying mesangium. In addition to its large perikaryon and the primary dendrites it originates the pPIN, differs from the other subtypes in that the axon generates a more elaborated FVC that distributes in the underlying blood vessel and its arising tributaries (Fig. 2h). Distally (i.e., beyond the vascular forking), the FVC reorganizes TFs (Fig 2h, asterisks) coursing along the vascular shaft like those we describe here for the sPIN.

**Figure 1.**
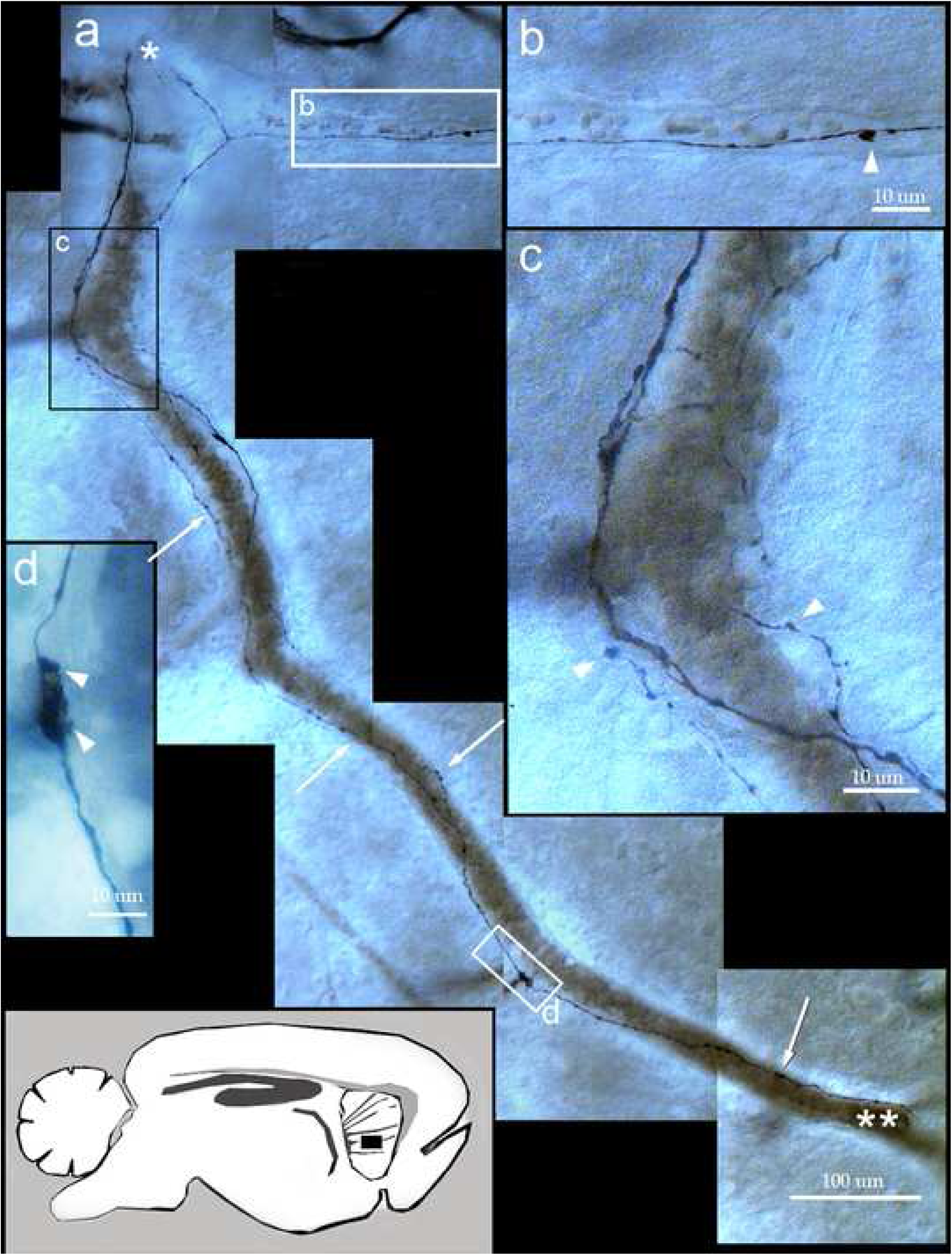
Parasagittal section of the rat caudo-putamen showing perivascular interneurons and *twin*, helicoidal fibrils following a branch the medial striated artery (extreme upper left). **a.** Photomontage of the artery proceeding from the subventricular neuropil (asterisk) into the basal ganglia. A perivascular neuron (framed “d”) issues long ascending fibrils that travel helicoidally about the blood vessel (arrows), from distal (double asterisk) to superficial (asterisk). **b.** High magnification view from the collateral boxed in “a”. The alternating, varicose appearance of the collateral fibril is interrupted by a balloon-like protrusion (arrowhead). Note the underlying capillary blood vessel harboring brownish, stacked red blood cells. **c.** Ramification of the blood vessel coexists with collateralization of the paired axons originating rounded outgrowths of assorted sizes stung by thinner fibrils (arrowheads). **d.** Soma and proximal processes of a spindle-shape neuron issuing divergent processes. As is shown in “a” the ascending axon provides daughter, twin fibrils, that having originated the plexus enlarged in “c”, resolve at the root of the blood vessel (asterisk). Adult rat brain, rapid-Golgi technique. Arrow heads = astroglial processes. Scale bars = 100 µm in a, 20 in b-d.

**Figure 2.**
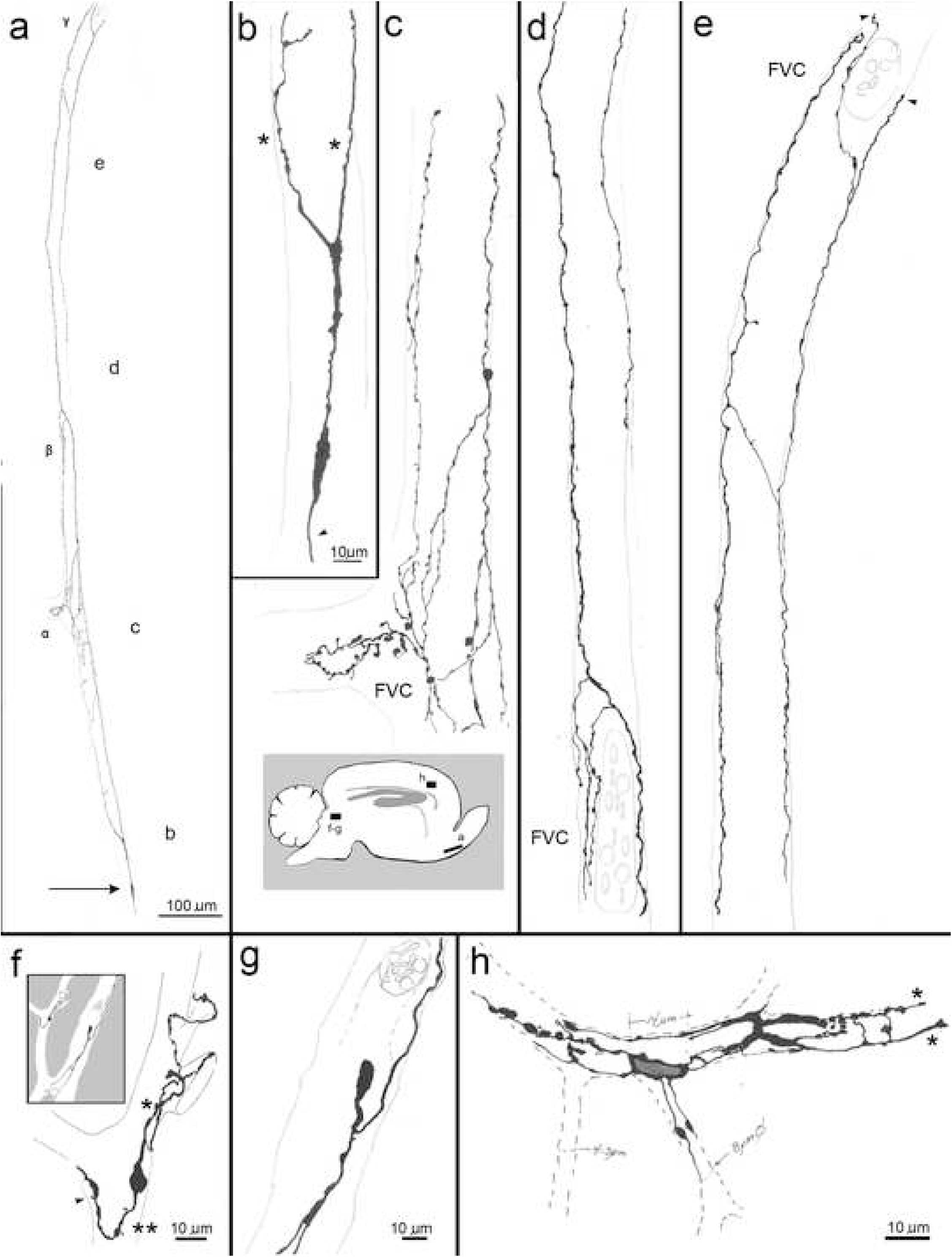
Camera lucida drawings of perivascular interneurons (PIN) and their processes in the adult rabbit brain. **a.** Spindle-shaped neuron (arrow) whose axon parallels a long, horizontal vessel traversing the anterior olfactory nucleus. α, β, and γ designate successive sites of the blood vessel ramification. To highlight is the position of the cell body at the distal part of the blood vessel, so that the axon and ascends against the blood flow which presumptively courses inversely from top to bottom. **b.** Enlargement displaying fine structural features of the spindle cell perikaryon with divergent proximal axon providing twin collaterals (asterisks) and a single dendrite (arrowhead). **c, d,** and **e** are sequential enlargements from the full axon shown in “a”. Noteworthy is that as soon as twin fibrils reach sites of vascular ramification, fibro-vesicular complexes (FVC) are formed. **f.** a bipolar type of PIN whose axon (asterisk) organizes a FVC at the root of an adjacent blood vessel. The neuron delivers a short, varicose dendrite (double asterisk) with sparse spines (arrowhead). **g.** Pseudomonopolar PIN. Notice that the axon gives rise to a fibro-vesicular complex which surrounds the vascular bifurcation, whereas the dendrite diverges to branch shortly afterwards. **h.** An stellate-shaped perivascular neuron. The short, varicose dendrites (left) with sparse spines, and the FVC (right side) converging into twin fibrils (asterisks) are illustrated. Scale bar in b applies to c-e.

TFs organize FVCs. TF lengths vary extensively but TF fibrils can still be seen in six- or seven-order ramifications in the gray matter of the cerebral isocortex, basal ganglia, and anterior olfactory nucleus, or elsewhere until the capillary diameter decreases <15 microns in the thalamus or brain -stem (Online Resource 1, 2) (1). An important remark that applies to the PIN and all its processes is that they lie embedded in EF (vide infra). As shown in a video (Online Resource 5), this location becomes obvious when the specimen is in and out of focus revealing that the distribution of the PIN processes match that of the astrocyte’s EF. Presumptive motor fibers identified in pial and parenchymal blood vessels display a relatively monotonous structure. They are composed small to medium sized vesicles stung by a thin fibril and, unlike the PIN processes, lie within the wall of the blood vessel (Fig. 3c, d).

**Figure 3.**
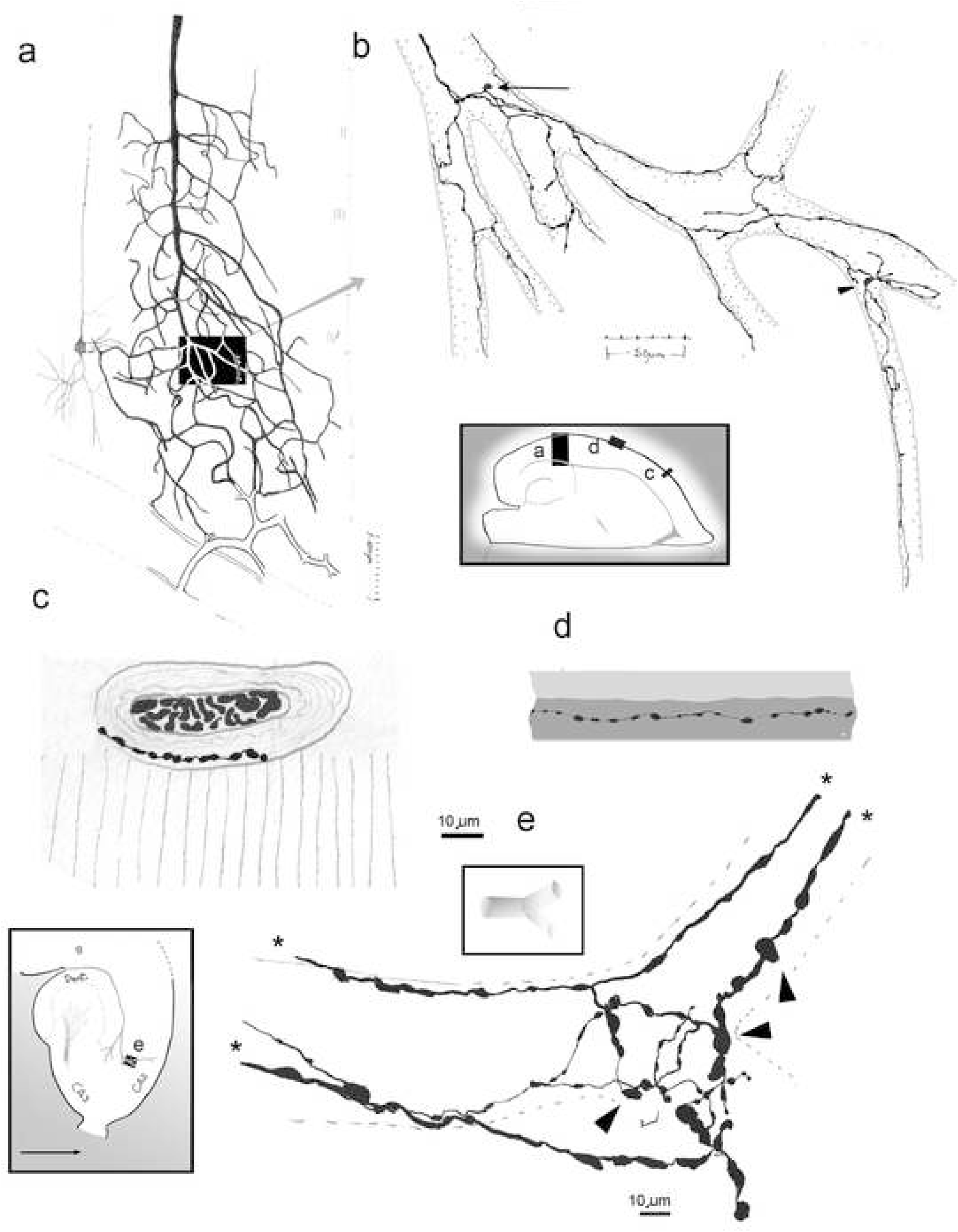
Camera lucida drawings illustrating perivascular interneurons (PIN) and axonal fibro-vesicular complexes in the cerebral cortex. **a.** Survey view of a long cortical blood vessel, tributary branches, and deep cortical veins (outlined in soft pencil) in the adult rat parietal cortex. Cortical layers are designated with Roman numerals in soft pencil. **b.** Higher view of the area boxed in “a”. Pattern of ramification and distribution of a pseudomonopolar (arrow) and a bipolar (arrowhead) PINs and their processes in the deep cortical blood vessels (soft pencil). Noticed the distribution of fibro-vesicular complexes associated with the blood vessel ramifications, and that thinner fibrils persist in small caliber blood vessels (low left and right sides). Rat parietal cortex. **c.** Arteriole at the brain surface (vertical lines in soft pencil). To note is a putative motor axon lies in the media between the smooth muscle fibers. **d.** Longitudinal section trough a meningeal arteriole with a putative motor ending distributed in the muscular wall (deep gray). Intima = pale gray. **e.**Horizontal section through a penetrating blood vessel proceeding between the rabbit subiculum (s) and dentate gyrus (dent.) to distribute in the hippocampus (i.e., CA2 and CA3 = Ammon’s horn, sectors 2 and 3) (inset). A fibro vesicular complex arises from in passage twin nerves (asterisks) about a site of vascular ramification (dashed). To note is the thin fibers alternating with bulb-like enlargements of assorted diameters (arrowheads). c-e rabbit hippocampus. Rapid-Golgi technique.

(1) Visualization of the long (>1 millimeter) axon and twin fibrils of a PIN-FVC requires immersion-oil (40 to 100x) observations and a thorough sampling of the brain (see Larriva-Sahd, 2008).

Implicitly a TF is composed of two unmyelinated fibers that run separately but have dissimilar calibers. In deep blood vessels the thicker fibril measures 0.5 to 1.2 µm in diameter, whereas its partner fibril is 0.1 to 0.4 µm in diameter. Both fibrils become progressively thinner as the underlying blood vessel proceeds towards the surface of the brain or brainstem parenchyma. However, 100 to 200 microns before resolving at the pia-brain interphase, TFs increase progressively in diameter to resolve as a discreet enlargement (Fig. 1a, asterisk, 2e, arrowheads). As shown in the panels of Figures 1a, c, 2b, e, TFs have no uniform diameters because distinct enlargements alternate with the shaft of each fibril. A peculiar feature of TF is that, as soon as the companion artery (see Marin-Padilla M 2012) ramifies, each fibril divides into two or three branches that anastomose and surround the roots of daughter vessels, thus organizing an FVC (Figs. 1c, 2c-e. 3e, 4). Aside from the marked pleomorphism of the FVC, PVBs and flat membranous structures are readily identified in order of frequency. As the observer proceeds distally from the vascular forking, fibrils of the FVC converge into a new set of paired, thinner TFs that run until the blood vessel ramifies once again (Figs. 1, 2a-e). This TF-FVC alternating pattern persists in successive ramifications and ceases when the capillary blood vessel diameter narrows <15 microns. Occasional FVCs may be also seen along the vascular shaft (Online Resource 2c). Species differences are noted. A first peculiarity is that the rabbit’s TF may undergo fasciculation so that thin (<1um) tightly packaged fibrils encompassing each TF can be seen at the highest magnification (Figs. 4e, f); this is especially noticeable just before parent TF originate a FVC. A second obvious difference identified in the rabbit’s FVC is its overall higher complexity as it exhibits many fibrils and large PVBs. The complexity of the rabbit’s FVC gives the impression that the rat’s FVC is the *trimmed* version (Figs. 3b, 4a-d) of the rabbit’s FVC (Figs. 3e, 4e, f). Lastly is the significantly higher number of assays required to obtain satisfactory FVC’s impregnation in the rat brain.

**Figure 4.**
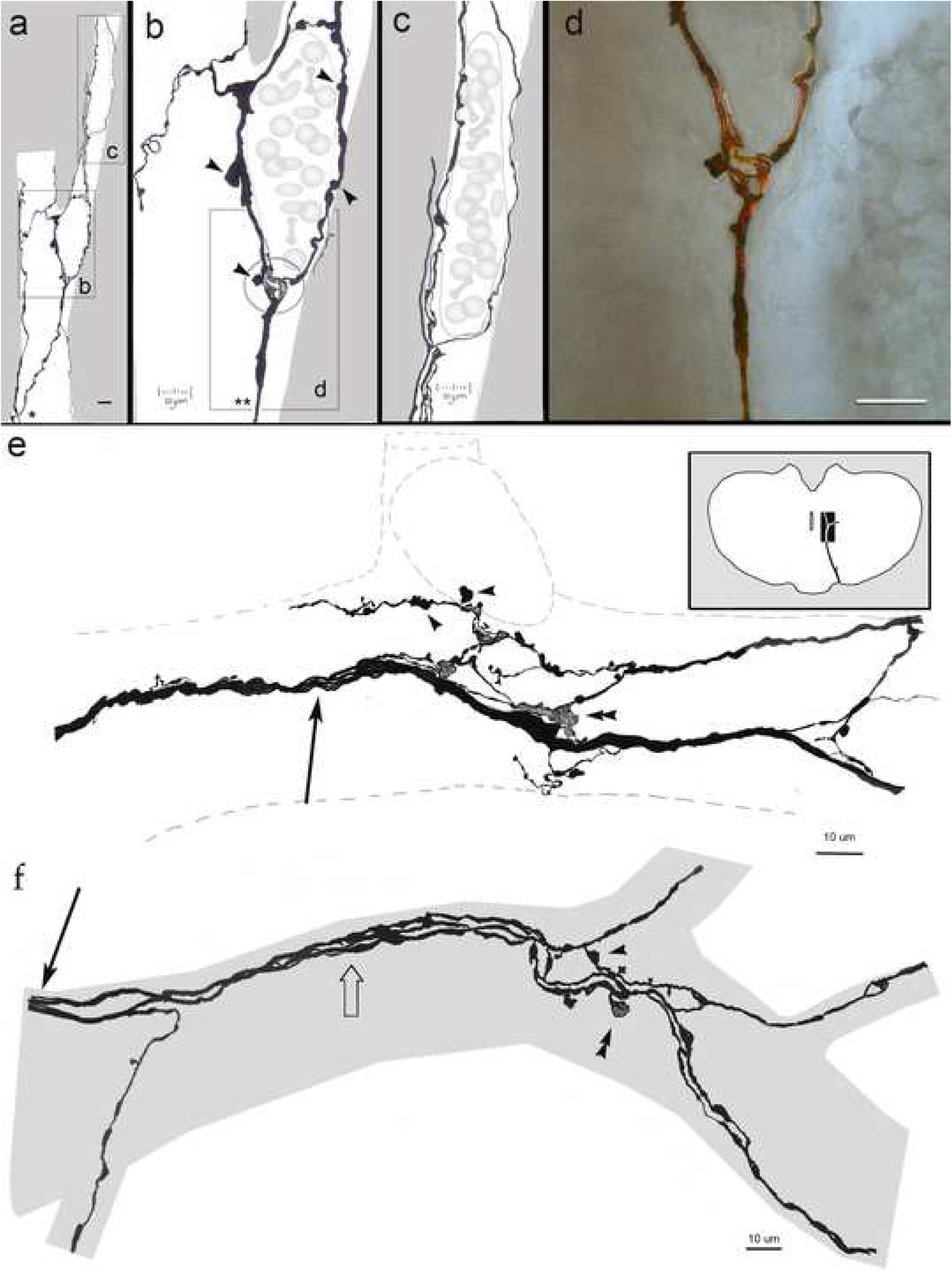
Images depicting the distribution of fibro-vesicular complexes associated with ramifying blood vessels as observed with the Golgi technique. **a.** Progression of twin nerves (asterisk) originating fibro-vesicular complexes at sites of blood vessel ramification. **b.** Enlargement from the area outlined in “a”. Ramification of a parent fiber (double asterisks) originating a series of flat (encircled) outgrowths alternating with bulb-like (arrowheads) formations. **c.** Enlargement from the squared area outlined in “a”, showing another fibro-vesicular complex. **d.** Photomontage from the area outlined in “b”. Adult rat hippocampus in a-d. **e.** Fibro-vesicular complex associated with a trifurcating blood vessel (dashed). Notice the fasciculation of the parent fiber (arrow) prior to giving rise to a fibro-vesicular complex. Arrowheads = bulbi. **f.** Fasciculated fibers (arrow) originating a fibro-vesicular complex. Arrowhead = bulb-like; double arrowhead = flat, membranous outgrowth. Adult rat a-d; adult rabbit e and f.

In the previous description illustrations from the frontal, parietal and piriform cortices, as well as basal ganglia, thalamus and midbrain have been used to highlight PIN and its processes. Demonstrative drawings of medulla oblongata and cervical spinal cord are available in the Online Resources 1 and 2, respectively. Searching for PINs in the cerebellar cortex was unsuccessful.

### 3.2 Histochemistry and Immunohistochemistry

Because of the different structure and topography between the TF-FVCs and the presumptive motor fibers and nerve endings (Fig. 3c, d) associated with cerebral blood vessels (Hartman et al. 1972; Cohen et al. 1997; Cauli et al. 2004; Hamel 2004; Cubelos et al. 2005), we hypothesized that they may correspond to a sensory ending (see Kruger et al. 2003; and Bewick 2015). Hence, primary antibodies directed against CGRP or VGlu 1, which are expressed and/or contained by sensory fibers-terminals and mechanoreceptors (Uddman et al. 1985; Ishida-Yamamoto et al. 1988; Silverman and Kruger 1990; Álvarez et al. 1993; Johansson et al. 1999; Fujiyama et al. 2001; Nakajima et al. 2007; Lennerz et al. 2008; Mazone and McGovern 2008; Bewick et al. 2015), as well as IB4 histochemistry, should reveal putative PIN-PVCs.

Detailed accounts of the CGRP-VGlu1-immunoreactive sites (CGRP-I and VGlu1-I, respectively) are available elsewhere (Efthekhari and Edvinsson 2011); hence, emphasis is given presently to immunoreactive structures associated with central blood vessels. In specimens incubated with antisera to CGRP and VGlu1, and histochemistry to IB4, horse-shoe and annular structures surrounding putative arteries and capillaries were visualized (Fig. 5a,b). In the eGFP mouse series of astrocytic EF outline and/or co-fluoresce with perivascular annuli. High magnification views (Fig. 5c) reveal small clusters positive to IB4-p with areas of CGRP-I and VGlu1-I co-labeling and overlapping. Again, perivascular eGFP end-feet next to the capillary wall surround and entangle the afore-mentioned clusters (Fig. 5b, c). Since nitric oxide (NO)-positive neurons have been implicated in mechano-sensory transduction (vide supra) and in influencing FH (see Idecola and Nedergaard 2007), immunohistochemistry and histochemistry were performed separately to visualize nitrergic neurons and their processes. Presumptive PINs laying on the capillary mesangium display strong cytoplasmic positivity to the NO synthesizing enzyme (i.e. NADPH) (Fig. 5d). Histochemistry to nNOS revealed thin, varicose fibrils outlining and/or encircling blood vessels (Fig. 5e, f).

**Figure 5.**
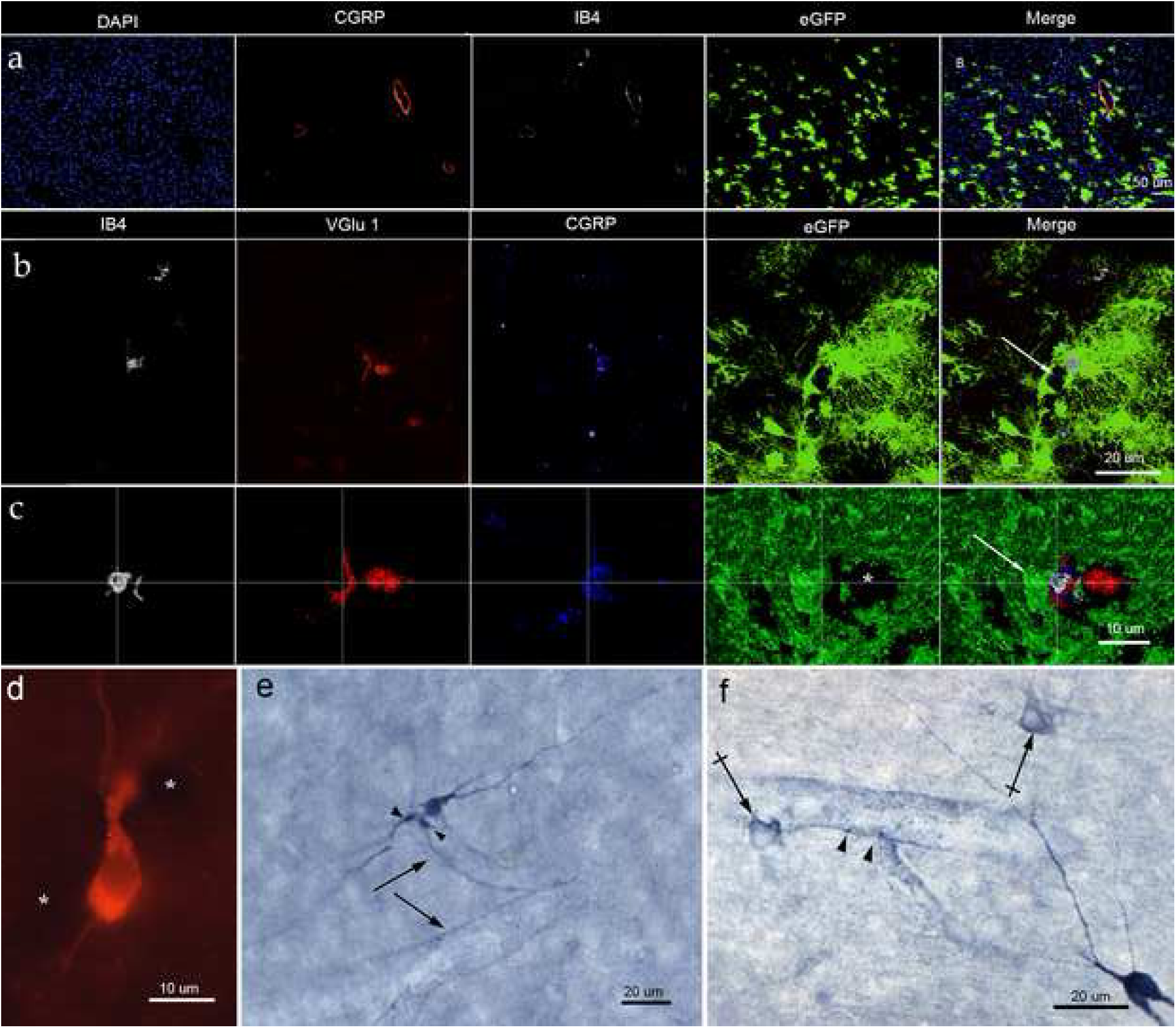
Fluorescent and confocal microscopy of cellular structures associated with central blood vessels. **a.** Survey picture from the posterior thalamic area. To note is the CGRP and IB4 fluorescence to donut-like structures outlining immuno-negative blood vessels with focal overlapping to eGFP of presumptive astrocytic end-feet. **b.** To note is the overlapping focal reactivity to IB4, with CGRP, and eGFP fluorescence alongside the lumen of a presumptive blood vessel (arrow). Parietal cortex. **c.** Immunofluorescence to a putative perivascular bulb encased by an end-foot (arrow) next to a blood vessel (asterisk). Amira software. Frontal cortex **d.** A perivascular nNOS positive neuron bounded by two blood vessels (asterisks). **e.** Neurons and processes associated to blood vessels. Notice the small, rounded, structures (arrowheads) beside a strongly positive NADPH neuron. Blood vessels appear outlined by thin, faintly positive, fibrils **f.** Ring-shaped processes (crossed arrows) and puncta (arrowheads) outlining the blood vessel’s wall. Parietal cortex in d – f.

### 3.3 Electron microscopy

As the fine structure of central neural and glial elements associated with the vascular wall has been extensively analyzed elsewhere (Jones 1970; Peters et al. 1976; McDonald 1983; Hamel 2004; Whitman et al. 2010) our description is addressed to the blood vessel wall in specimens from the frontal and parietal isocortex, posterior thalamus, and olfactory bulb medulla. Due to the superior fixation accomplished in the rat specimens, 3D reconstructions were made in this species.

PINs. Survey views of the putative PIN’s perikaryon reveals its slender, cigar-shaped contour (Fig. 6a, g). The scarce cytoplasm averages a 2:10 cytoplasm/nucleus ratio, it contains poorly differentiated organelles including abundant free ribosomes and scattered mitochondria; cisternae of the rough endoplasmic reticulum are indistinct (Fig. 6a, b). A solitary dendrite coursing parallel to the capillary wall is commonly observed. In 3D reconstructions performed in two neurons of the cerebral cortex, the origin of the axon was fully visualized. In one reconstruction, the axon arises from the lateral surface of the perikaryon (Fig 6 e, g) and in the other one (not shown), from the cell pole opposite to the dendritic process. As featured elsewhere for neurons of the bed nuclei of the stria terminalis (Larriva-Sahd 2006), the initial segment of the axon contains clusters of small, clear- and dense -cored -vesicles, both entangled by neurotubules. The bounding plasma membrane of both the initial segment (Fig. 6e) and the distal axon (Fig. 6f) is underscored by a stripe of fuzzy, electron-dense material. The cell body and proximal processes are surrounded by the capillary basal lamina and neighboring neuropil, whereas the axon courses underneath or surrounded by EF (Fig. 6f). One or two synaptic boutons define asymmetrical contacts with the PIN perikaryon (Fig. 6a, d), resembling those described by Blasko et al. (2013) for nNOS-positive neurons in the rostral migratory stream.

**Figure 6.**
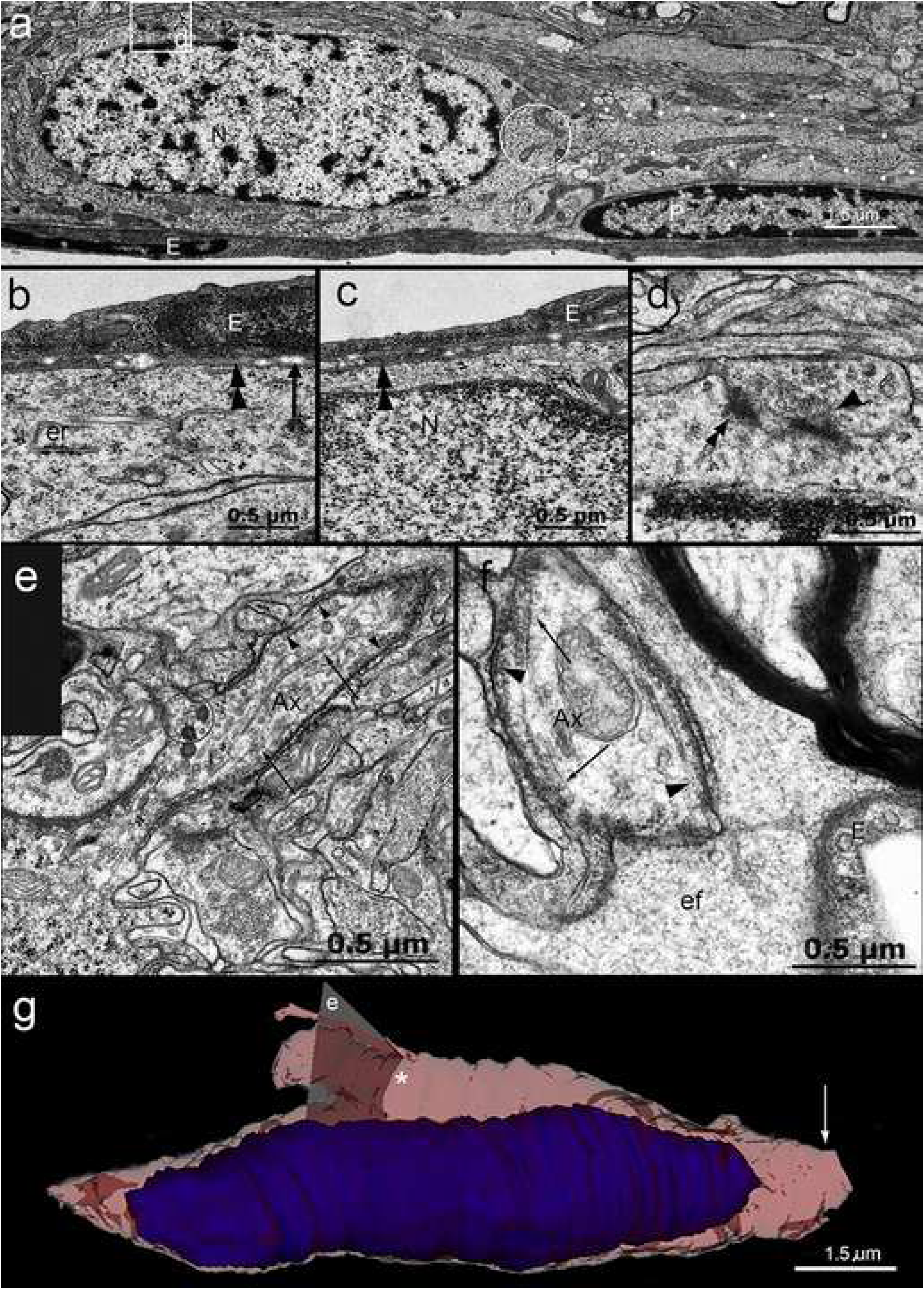
Inner and outer structure of the soma and proximal processes of perivascular interneurons in the cerebral cortex. **a.** Bipolar, possibly, spindle-shaped neuron paralleling a capillary blood vessel. The neuron’s cytoplasm is scarce, containing numerous free ribosomes and scattered mitochondria (circle). A thick, solitary dendrite (dots) courses horizontally. E = endothelial cell, P = pericyte. **b.** Direct interaction of the dendrite (center) of a perivascular neuron from the hippocampus. The process lies in direct apposition with the basal lamina (double arrow-head) that embeds collagen fibrils (arrow). er = endoplasmic reticulum. **c.** Perikaryon of a perivascular neuron in contact with the capillary basal lamina (double arrow head) surrounding the overlaying endothelium (E). N = cell nucleus. **d.** Asymmetrical, axo-somatic synapse in the neuron shown in “a”. Notice are the numerous rounded vesicles (arrowhead) polarized toward the synaptic active zone (double arrow head). **e.** Axon initial segment of a perivascular neuron. The axon arises from the soma (bottom left side) and contains small bundles of neurotubules (arrows) and assorted vesicles (circle). To note is the fussy, electron-opaque material (arrow-heads) underscoring the axon (Ax) plasma membrane. **f**. unmyelinated axon (Ax) from perivascular neuron encased by an astrocytic end-foot (ef) next to a capillary blood vessel. E = endothelial cell cytoplasm. Arrows = neurotubules, arrowheads = submembranous electron-dense material. **g**. Reconstruction of the perivascular neuron perikaryon. Notice the scarce cytoplasm (pink) surrounding the cell nucleus (blue), the sidewise initial segment of the axon shown in “e” (asterisk), and the root (arrow) of a dendrite. Series of two hundred and twenty sections assembled with the Reconstruct software.

Observations directed at the perivascular and mesangial neuropil reveal the presence of groups of bulb-like structures matching in size, shape, and location (Figs. 7-10) to those of the FVCs as observed at the photic and fluorescence light microscopes. PVBs organize clusters of two to five and lying between the capillary endothelium (McDonald 1983; Bruns and Palade 1968; Jones 1970 Peters et al. 1976) and the surrounding neuropil. There, PVBs are in apposition to-, or embedded in the endothelial and pericyte capillary basal laminae and, EF (7a and b, 8a, d, f, g), and are in focal contact with Mato cells (Mato et al. 1984)(not shown). Observations of a series of up to one-hundred fifty sections reveal that each PVB derives from an unmyelinated axon, as suggested by single sections (Fig. 6e, f). Figures 7a, c, d, and 10 show a narrow *proximal* axon that enlarges abruptly to the PVB itself, which narrows in the opposite pole to structure a *distal* axon, with the same inner structure than the former. Terminal PVBs arising from a stem axon, although infrequent, are revealed by 3D reconstructions (Fig. 10c).

**Figure 7.**
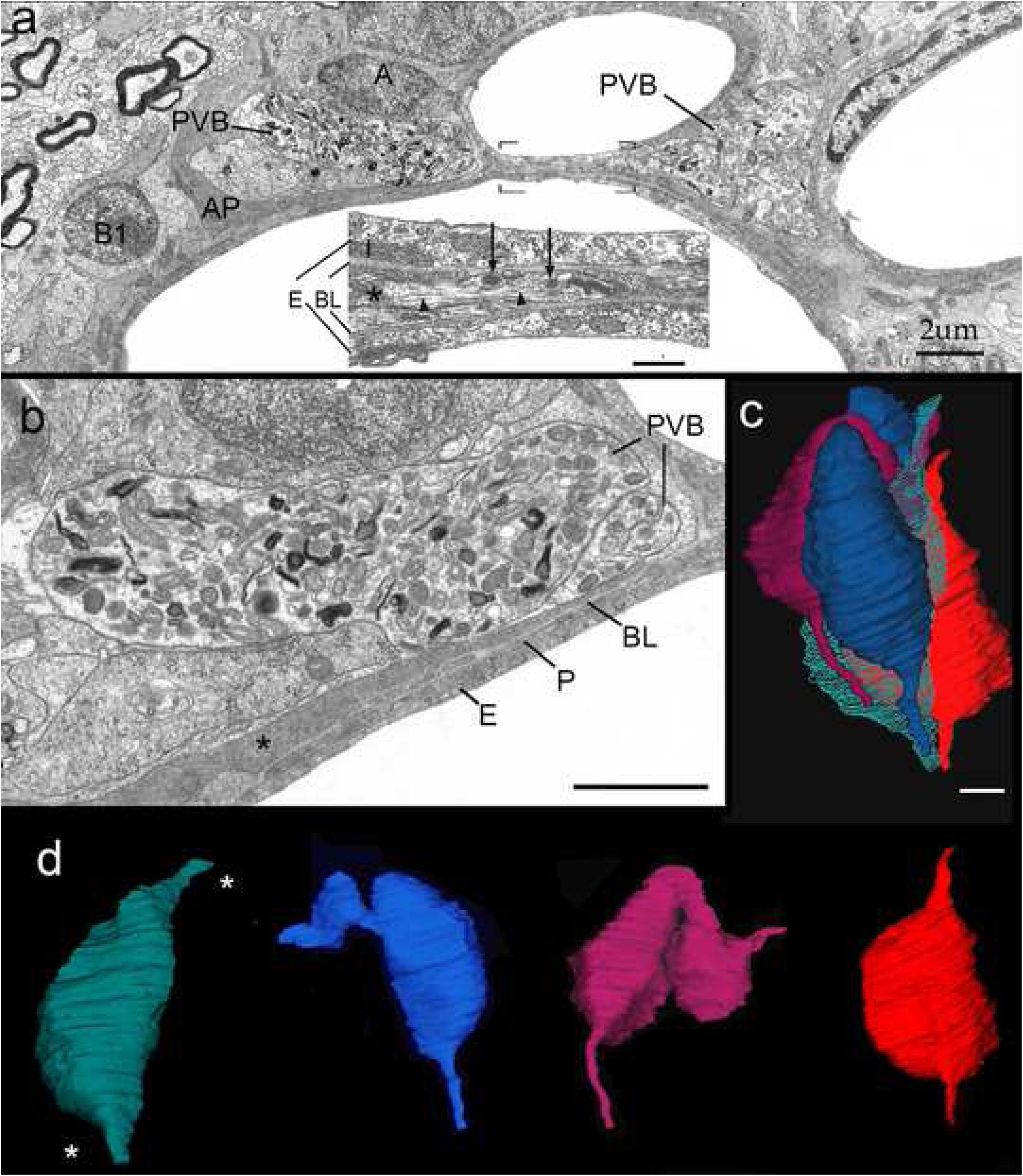
Bi- and 3-dimensional electron microscopic views of clusters of perivascular bulbs (PVB) contiguous to a presumptive vascular trifurcation within the rat olfactory bulb medulla. **a.** Survey micrography showing the interactions of PVBs with capillary basal laminae, astrocytic processes (AP), and adult-born cell-precursors (A and B1) (Doetsch et al. 1997). **i.** High magnification depicting an axon (asterisk) arising from a PVB which passes between the two capillary basal laminae (BL) underscoring endothelial cells (E). The PVB axon contains dense-cored vesicles (arrows) and sparse microtubules (arrowheads). **b.** PVBs bounded by two capillary blood vessels. The numerous mitochondria and electron-opaque membranous organelles contained by a limiting cell membrane of smooth contour are evident. Asterisk = process of a putative A cell (see above), E = endothelium, P = pericyte, and BL = basal lamina. **c.** Reconstruct of a cluster of four complete PVBs contained in one-hundred and eighty-four consecutive sections. **d.** Outer aspect of the bulbi shown in “**c**”. Noticed is that each bulb results from the expansion of an axon that narrows at the poles (see “a, inset”) process proceeding at either bulbar pole (asterisks).

The PVB itself contains assorted membranous organelles entangled by small bundles of neurotubules. The organelles include numerous, laminated secretory granules (SGs), mitochondria and multivesicular bodies, in decreasing frequency (Fig. 8). SGs contain concentric membranes embedded in an electron-dense matrix and although generally rounded, the presence of flat, elliptical, or pleiomorphic shapes are, nonetheless, common (Fig. 8). Frequent fusion of SGs with the limiting plasma membrane and an occasional reinforcement of its outer aspect is observed (Figs. 8b, between arrows, c). Overall, the organelles described here for the PVB, and, especially its SGs, recapitulate those observed in the Meissner corpuscle from the same animal, Online Resource 3, and elsewhere (Hashimoto, 1973 Ide et al 1987), as well as in sensory endings of the Pacinian corpuscle and other peripheral sensory receptors (Chourchkov 1973; Kimani 1992; Malinovsky 1996; Maeda et al. 1999; Iigima and Zhang 2002; Kruger et al. 2003).

**Figure 8.**
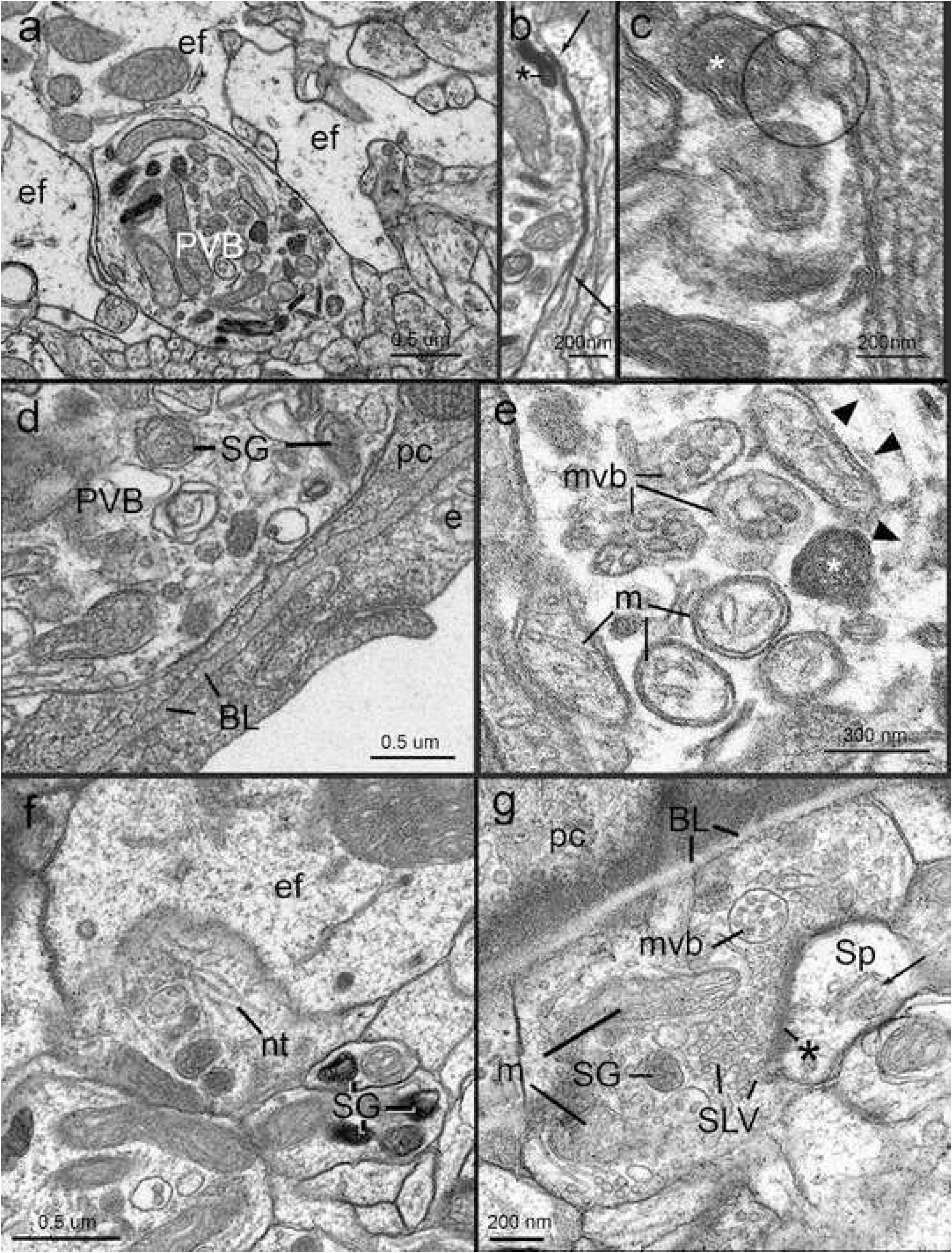
Organelles and interactions of perivascular bulbs (PVB) within the neurovascular unit of the adult rat. **a.** A PVB partially encased by astrocytic end-feet (ef) and neuropil (bottom) which contains laminated secretory granules, mitochondria and assorted vesicles. **b.** Outer aspect of a PVB out-lined by an astrocytic process. Notice the fusion of a laminated secretory granule (*) with the bulbar plasma membrane which is reinforced by an electron-opaque material (between arrows). **c.** Fusion of a membranous secretory granule (asterisk) with the plasma membrane (circle). c-d = posterior thalamus **d.** Relationships of a PVB with the capillary wall. To note is that the PVB faces directly with a pericyte (pc) and endothelial basal laminae (BL). e = endothelial cell. **e.** High magnification micrograph of a PVB containing multivesicular bodies (mvb) bounded by mitochondria with vesicular cristae (m), neurotubules (arrowheads), and a laminated secretory granule (asterisk). d-e = Frontal cortex **f.** Custer of PVB, one of them, facing end feet (ef). SG = laminated secretory granules, nt = neurotubules. Parietal isocortex. **g.** A PVB partially covered by the basal lamina (BL) of a pericyte (pc). To note are the numerous synaptic vesicles (SLV) clustered next to an asymmetrical synaptic interaction with a dendritic spine (Sp). Arrow = spine apparatus; asterisk = active zone; m = mitochondria; SG = secretory granule. Parietal cortex.

Another specialization that is observed in about one third of PVBs in the neocortex, hippocampus, and posterior thalamus, consists of typical asymmetrical synapses (Fig. 8f, g, 9a-c) (Peters et al. 1976). These PVB synapses consist of collections of small, rounded-synaptic-like vesicles (SLVs) with electron-lucid matrices that converge onto a synaptic contact shared with a dendritic spine. Small membranous collections forming a postsynaptic specialization are frequently seen within the cytoplasm of the spine itself (Fig. 8g). Usually the PVB-synapse is partly or completely surrounded by EF (Figs. 9a-c). In 3-D reconstructions it is noted that SLVs are polarized towards the synaptic active zone whereas SGs distribute throughout the PVB cytoplasm (Fig. Fig. 9d, e).

**Figure 9.**
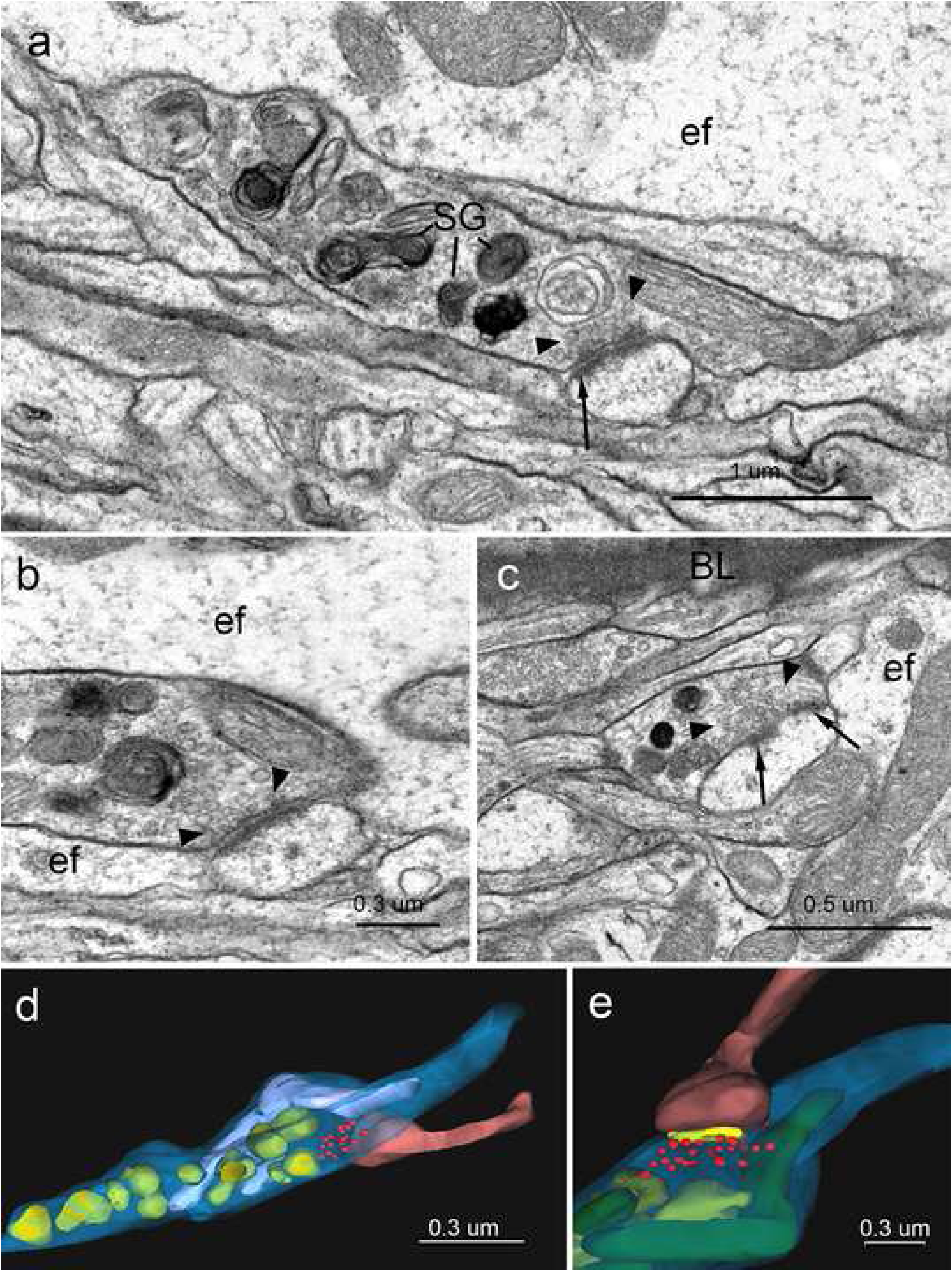
Synaptic interactions of perivascular bulbs (PVBs). **a.** A perivascular bulb forms an asymmetrical synapse with a nearby spine in the underlaying neuropil. To note is the cluster of small clear, presynaptic vesicles next to an active zone (arrow) distributed along the presynaptic cytoplasm. SG = laminated secretory granules. Mesencephalon. **b.** High magnification view of a PVB making an asymmetrical spinous synapse encased by astrocytic end-feet organizing a tripartite synapse (Araque et al 1999). Frontal cortex. **c.** PVB-spinous asymmetrical contact with two active zones (arrows). Arrowhead = synaptic vesicles; BL = capillary basal lamina; ef = end foot. Deep parietal cortex. **d.** 3-D reconstruction of the PVB shown in “a”. The PVB cytoplasm (blue) harbors numerous laminated secretory granules (green), two mitochondria (light blue) and a cluster of synaptic vesicles (red) underneath the site of apposition with the dendritic spine (pink). **e.** Close-up of the synaptic interaction shown in “**d**” to which the active zone (yellow) has been added. Reconstruct software.

PVBs are housed in a complex framework formed by the elements of the NVU. Both the PVB and its parent axons are enveloped by an elaborate cuff composed of the capillary basal laminae and astrocytic EF in variable proportions (Figs. 9a-c, 10). An exception for this is noticeable in capillaries of the olfactory bulb medulla whose neuronal precursors and their processes envelop a substantial part of the PVB surface (Fig.7a, b). Aside from this, PVBs are invariably embedded in the basal laminae, which organizes an amorphous receptacle of variable thickness. PVB areas devoid of basal membrane are covered by EF, thereby providing a complete PVB encasing. In addition to the glial-basal laminae interaction, the surface of a small subset of PVBs in the olfactory bulb and cerebral cortex is directly exposed to the neighboring neuropil (see above) (Fig. 8a).

**Figure 10.**
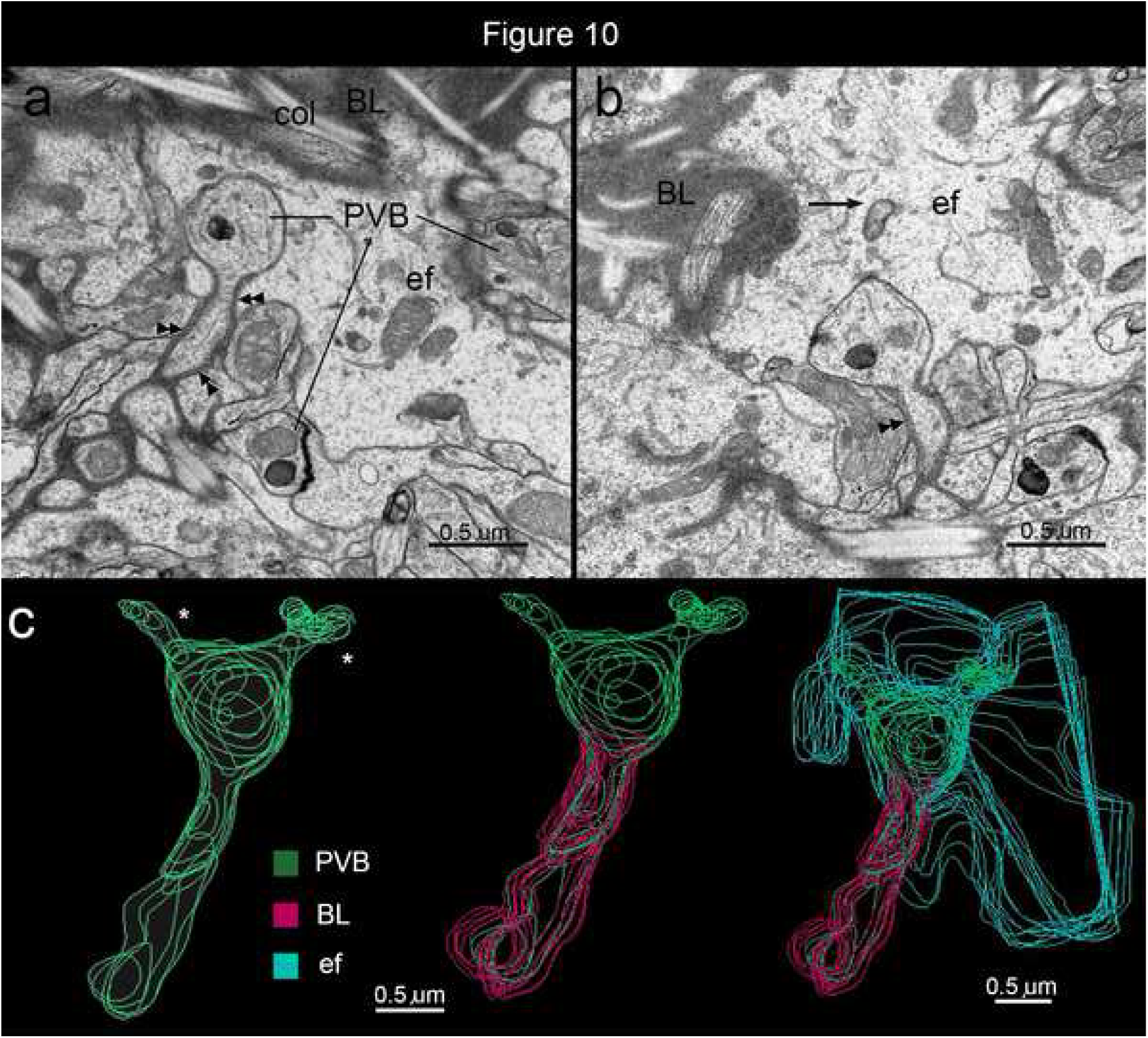
Series of electron micrographs depicting the structure and interactions of perivascular bulbs (PVBs) with the neurovascular unit. **a.** The tangential section through the periphery of a capillary blood vessel exhibiting three PVBs. Notice that the bulb on the left is immersed by astrocytic end-feet (ef) whereas basal laminae (arrowheads) covers its stem axon. BL = capillary basal lamina; col = collagen bundles. **b.** Specimen fourteen sections apart from that section shown in “**a**”. Notice that the bulb and its stem fiber remain encased by the ef and basal lamina (arrowheads), respectively. Arrow = bulbar appendage. **c.** Stepwise reconstruction from the bulb and its stem axon displaying the covering built-up by the basal lamina (red) and astrocytic ef (turquois). Notice the bulbar paired appendages climbing toward the blood vessel wall (not shown). Frontal isocortex, adult rabbit.

## 4. Discussion

### 4.1 Perivascular interneuron or ganglion cell?

We describe a unique group of perivascular interneurons or PINs intrinsic to the NVU of the adult brain and brainstem vasculature. Conservation of them was found in a lissencephalic (i.e., rat) and a gyrencephalic (i.e., rabbit) species. For its outstanding distribution and interactions with the NVU, the PINs should be considered a new cell type. Overall is the PIN’s soma and its processes that are confined to the perivascular domain, frequently surrounded by the capillary basal laminae and astrocytic EF. Although neurons sharing the PIN’s perivascular location have been previously described, to the best of our knowledge, information on distribution and organization of their processes was not available. Hence, we opted for the Golgi technique that has been effective in defining most neuron types and peripheral receptors within the nervous system. The PIN’s axon issues clusters of terminals, termed here PVBs whose morphology, reactivity, and immunoreactivity match with those observed here and elsewhere in the Meissner corpuscle and in other peripheral mechanoreceptors. From its cytological characteristics the PIN may be considered an elementary ganglion cell (GC). In fact, like the peripheral branch of a GC, the PIN’s axon gives rise to a presumptive sensory ending. Furthermore, glial and perivascular basal laminae observed in mechanoreceptors (Andres and During 1973; Dubvový and Bednárová 1999) encase the PIN’s putative sensory ending. Even more, the TF fasciculation and sequential distribution of the sPIN-FVC axon at sites of vascular ramification might be regarded as a central form of functional segmentation. The early assumption of the peripheral GC as a mere relay neuron proved to be an oversimplification; since it receives several peripheral inputs and its central process branches before entering to the spinal cord (Langford and Coggeshall 1981). Hence, spatially dissimilar, segmental receptor fields recruit the same GC. This soon afterwards confirmed by double labeling from different sensory fields (Christianson et al. 2007), supports the ability of the GC to decode time-space sensory cues. By the same token, the sPIN axon provides a series of PVCs that distribute sequentially in the arising capillary blood vessels thereby suggesting spatial decoding by the PIN itself (Figs 2a, 11). Given that the decreasing intensity of the blood flow as it travels along the blood vessel (Hall 2016), the PIN and its FVC series should be equally exposed to a decreasing dynamical force (Fig. 11d). This suggests that like a GC, the PIN gathers dissimilar segmental inputs (Fig. 11a, d) (see Larriva-Sahd 2014). While this hypothetical ability of the PIN to decode time-space inputs is open to future neurophysiological assessment, its striking analogy with the peripheral GC suggests that it corresponds to an *elementary ganglion cell*. Although drawbacks of adducing function from structure alone notwithstanding, novelty of this putative receptor admits some functional correlates as discussed next.

**Figure 11.**
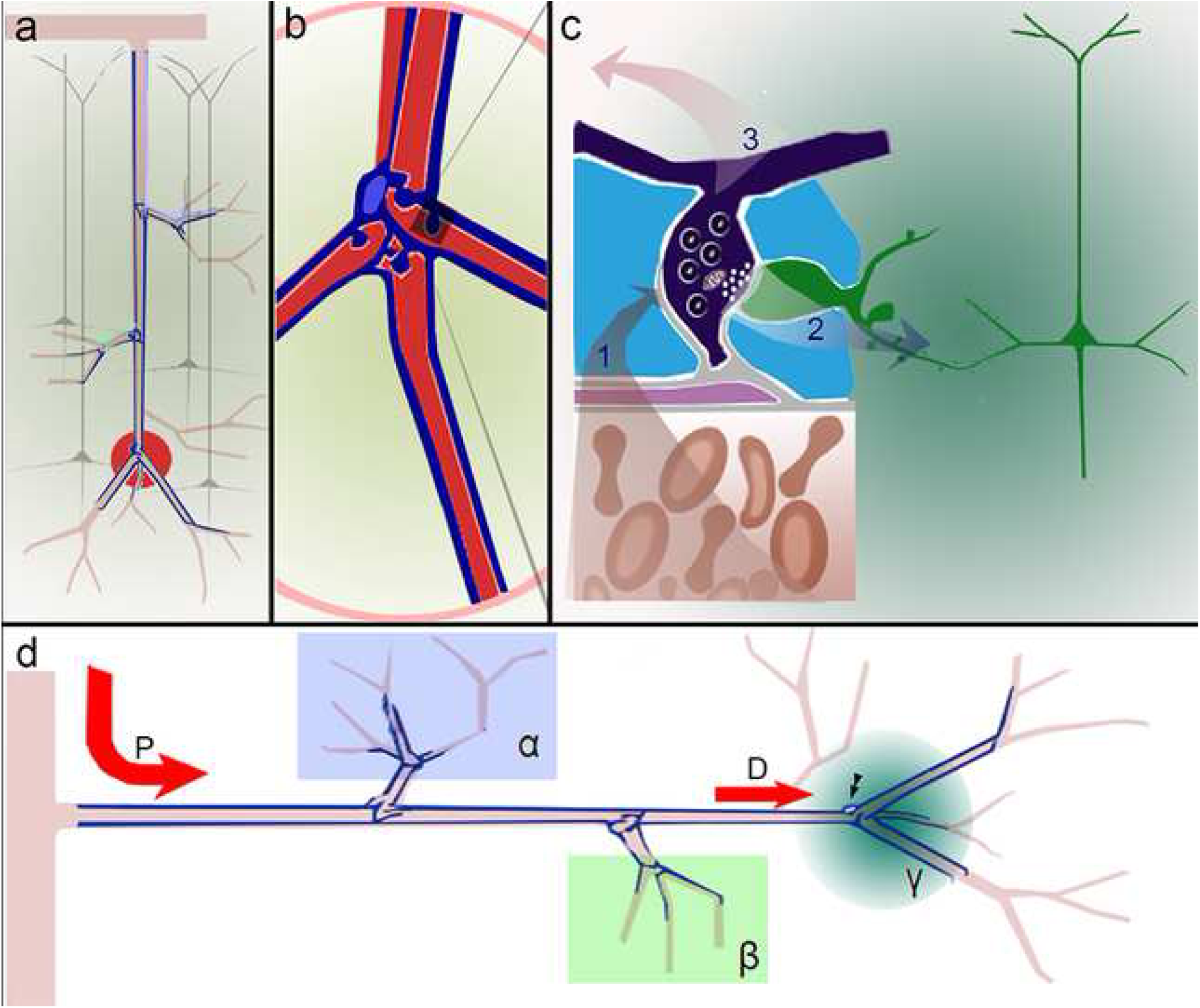
Cartoons illustrating morpho-functional correlates of perivascular interneurons (PIN), axons, and fibro-vesicular complexes (FVCs) they originate. **a.** A set of deep perivascular neurons gives rise to long, ascending axons that, provide series of fibro-vesicular specializations about the blood vessel ramification (one of them encircled in red). **b.** Within the neurovascular unit, each FVC originates numerous perivascular bulbs (one of them shaded). **c.** Putative feed-back control of blood flow. Blood flow (bottom) produces a mechanical impact in the neurovascular unit, including perivascular bulbs (deep blue) (Vector 1) leading to a receptor, i.e. graded, response carried out by the latter. Receptor potentials may travel to the cell body (vide infra) (Vector 3) or provoke neurotransmitter release on dendritic spines of nearby neurons (Vector 2). **d.** To note is that the sequential arrangement of FVCs from the proximal to the distal part of the artery may *sense* both local mechanical blood flow impact within each ramification and the overall blood flow within the PIN perikaryon (arrowheads). For instance, opening of the blood vessel in (α) during functional hyperaemia yields a decrement in the stem blood vessel (Vector D) which may, in turn, be sensed by the perikaryon and proximal processes of the PIN at the “γ” territory (turquoise). As implicit, an additional blood flow demand in territory beta (β), would lead to an inverse decrease in the overall magnitude of vector D corresponding to territory gamma (γ). If the PIN, like other neurons in the CNS (see Larriva-Sahd, 2014), decodes temporal and spatial inputs, it would then be able to respond to segmental blood flow variation (i.e., α, β, and γ) as well as to the resulting overall blood flow (vector D).

### 4.2 Perivascular neuron connectivity

While we were unable to find projecting axons from the PIN to the neighboring neuropil the synaptic interactions of PVCs with typical dendritic spines (Arellano et al. 2007) provide a valuable connectional clue. In fact, association of synaptic-like vesicles (SLV) with the neighboring glial and stromal cells has been thoroughly investigated so that consensus about their functional role has been reached (Bewick 2015). At first glance, the presence of SLVs within the primary sensory endings was found to be paradoxical, or at least heterodox (see Kruger et al. 2003). Subsequent experience supported the SLV involvement in vGlu 1-mediated glutamate recycling at the mechanosensory ending-receptor interphase (Bewick and Banks 2015) further suggesting that the SLV-receptor *contacts* is the structural signature for the sensory ending-mediated tune-up of the receptor itself (Bewick 2015). Reactivity to the vGlu1 antibody detected here in putative PVBs lends further support to its receptor nature. However, there is a fundamental difference between the PVB-synapse and the contact of the primary sensory nerve-receptor interaction. SLVs in sensory endings polarize towards the glial or cellular elements associated to the receptor; whereas SLVs in the PVB organize the presynaptic element of a typical axo-spinous synapse. Hence, the PVB appears to modulate nearby *neurons* via chemical synapses. How might sensory cues, mechanical or otherwise, elicit RPs in the PVB? Although this cannot be answered here, the similarities of the vascular and glial elements associated with the PVB with those of peripheral mechanoreceptors are striking.

### 4.3 Stromal and glial components of the perivascular bulb structure a perivascular organ (PVO)

Sensory endings and associated glial and stromal structures define sensory organs. The PVB and related endothelial cells, pericytes, EF, and extracellular matrix, appear to counterpart them. Although endothelial cells (Busse and Fleming 2003; Davis 1995) and astrocytes (Davis 1995; Filosa et al. 2016) are both necessary to trigger FH and both display varying degrees of response to mechanical stimuli, there is no evidence that they transduce them to RPs. Vascular and astroglial cells as well as the extracellular matrix are nonetheless crucial for sensory transduction (see Zimmerman et al. 2014; Nesslinger 1996). What might the involvement of each of them be? Exposition of the endothelial cell to the shear stress imprinted to the vascular wall leads to the synthesis and release of potent vasoactive molecules (Busse and Flemig 2003). Notably, nitric oxide and the endothelial-derived hyperpolarizing factor (McGuire et al. 2001) thought to modulate arterial smooth muscle tone. Complementary, astrocytes which form a bounding annulus around the endothelium are also necessary to trigger FH (Idecola 2004). At this point, the close interaction of the astrocytic EF with the PVB-synapse described here may be relevant. In fact, association of astrocytic processes enveloping both pre- and postsynaptic elements represent the structural substrate for the astrocyte’s modulatory influence on glutamate-mediated transmission (Araque et al. 1999; Tasker et al. 2012). Since astrocytes are highly responsive to blood flow variation (Idecola and Nedergaard 2007; Attwell et al. 2010; Filosa et al. 2016), it is further suggested that EF may modulate the eventual PVB synaptic outcome (Araque et al. 1999), likely to be vGlu1-mediated. An emerging corollary is that the PVB-spine synapse is a physiological link between the putative receptor described here and the neural circuitry involved in sensory decoding for an appropriate vasomotor response.

### 4.4 Sensory innervation of the brain vasculature: functional implications

Functional brain parceling is based upon variation in FH elicited by cellular energy demands. Notably, functional magnetic resonance, near infrared spectroscopy, and other non-invasive procedures take advantage of the dynamic, differential blood flow to define functional recruitment of discrete territories of the brain. (see Idecola 2004; Filosa et al. 2016). While in the systemic circulation the cardiac out-put is constantly monitored by specialized sensory organs, it is poorly understood how changes in blood flow during FH feed-back vasomotor responses. The PIN described here represents an attractive candidate which is prone to be explored. In fact, utilization of in vivo recordings combined with pharmacological tools should help in defining the physiological role of the neurochemicals identified here within in association with PVBs.

Given the paramount role played by blood vessels and supply in metabolism, nourishment, development, and plasticity of the normal and pathological brain, a systematical account reviewing all potential implications that mechanoreception might have, should be left as an open chapter (see Ward and Lemana 2004; Idecola 2004). However, the association of PVBs with cells and their processes phenotypically indistinguishable from those of adult-born neuron precursors observed in the olfactory bulb medulla (OBm) (Doetsch et al. 1997)(Fig. 7a, b), motivate some prompt remarks. The OB is one of the privileged brain areas of the adult brain in which new-born neurons migrate from the wall of the lateral ventricle to the bulbar cortex where they incorporate to preexisting neural circuits. Blood vessels and glial cells are intimately implicated in viability, induction, and guidance of prospective adult-born neurons (Whitman et al. 2010). It is admitted that blood vessels (Whitman et al. 2010), extracellular matrix (Hallmann et al. 2005), and astroglia (Lois and Alvarez-Buylla 1994) guide differentiating neurons to their eventual targets, a process involving numerous signaling and trophic molecules expressed and/or released by and within by the NVU itself (Ward and Lemana 2004). But how do prospective neurons recognize the source and the target for a successful journey? In effect, a blood vessel i.e., cylinder, lacks *per se* polarity to modulate a sequential expression and/or secretion of signaling molecules and trophic factors by the NVU itself. A possibility is that the putative receptor described here transduces *directionality*, i.e., that generated by the FH, to the former (Fig. 11d). It is tempting to imagine that the PIN and its stromal and glial partners would, directly or indirectly *instruct* neuron-precursors about eventual site(s) where increased metabolic demands require them (Sharp et al. 1975; Magavi et al. 2005, Alonso et al. 2008). Functionally recruited olfactory glomeruli trigger FH (see Xu et al. 2000; Alonso et al. 2008; Chaigneau et al. 2003) so that blood flow from tributary blood vessels is redirected to the former. Because of the negligible elasticity of the blood column a functional *sink* is created. Our observations in Golgi-impregnated specimens show that blood vessels in the OBm ascend radially to the olfactory cortex, resolving into glomerular veins and venous sinuses (Online Resource 4); accordingly, FH within a given glomerulus should modify blood flow retrogradely. Although arterial and venous blood vessels may be readily identified with the Golgi technique (Marin-Padilla 2012), it still be arguable that blood flow direction cannot be defined in a histological section. However, this does not challenge the essence of our postulation, whether the blood flows towards (See Lecoq, et al. 2009) or from a glomerulus it *impacts* retrogradely or anterogradely blood flow (Hall 2016), respectively. It is therefore speculated that decoding of blood flow variation by series of PVBs (Fig. 11d) would enable the NVU to conduct the migratory processes towards functionally active territories.

## 5. Concluding remarks

Lord Adrian (1954) described the physiological characteristics of mechanoreceptors and determined that stretching of tendinous sensory endings (Sherrington, 1892) results in RPs whose frequency varies as a function of the intensity of the stimulus. This remarkable property of receptors was soon extended and found to operate in all sensory organs and terminals. In the systemic blood flow, receptor organs and endings are strategically positioned to sense and transduce physic-chemical cues to RPs that are decoded by the CNS to perform appropriate motor responses by the heart and blood vessels (Hall 2016). The structural identification of a putative mechanoreceptor in precapillary and capillary blood vessels within the CNS opens the possibility that, like in the systemic blood circulation, subtle hydrodynamic variations of distal blood flow are likewise transduced to neuron chains for a successful adaptative motor outcome. The associations of PIN-PVCs with the NVU as a counterpart of the peripheral GC-receptor supports the existence of brain proprioception (i.e., self-sensorial).

## Acknowledgements

This work was supported by CONACyT, Grant 1782 to LC and JL-S and by Universidad Nacional Autónoma de México, PAPIIT Grant IG200117 to LC and JL-S. The transgenic hGFAP-GFP mouse line is a generous gift from Dr. Helmut Kettenmann. Authors appreciate the suggestions made by Dr. Carlos Cepeda to improve our manuscript and thank Gema Martínez-Cabrera, Carlos Lozano-Flores, Lourdes Palma, Elsa Nydia Hernández-Ríos, Martín García, Rafael Olivares for technical assistance. The through revision of our manuscript by Jessica González Norris is also appreciated.

## Online Resources

**1.** Camera lucida drawing showing the pattern of ramification of perivascular nerves surrounding a blood vessel piercing the dorsal horn (upper right). Adult Rabbit, cervical spinal cord.

**2.** Camera lucida drawings showing twin fibers and fibro-vesicular complexes associated with blood vessels in the brainstem. **a**. Low medulla oblongata. **b**. Upper medulla oblongata. c. Pons. A single fibril originating a fibro-vesicular complex along the shaft of a blood vessel. Adult Rabbit. that axons and fibro-vesicular complexes vesicular complexes lie among the astrocytic end-feet that form a shell (deep brown) about them.

**3. a.** Survey picture of the Meissner corpuscle of the adult rat palmar skin. A sensory ending (single asterisk) containing numerous mitochondria and sparse laminated secretory granules (arrow) is encased by the concentric glial lamellae (double asterisks). **b.** high magnification view of a sensory ending in a Meissner corpuscle. To note are the two secretory granules (arrows) having concentric, laminated cores. **c.** enlargement from a perivascular bulb at the same magnification containing a secretory granule (arrow) and a multivesicular body resembling those shown in the Meissner corpuscle (b).

**4.** Distribution of the blood vessels in the adult rat olfactory bulb. Sagittal views. **a.** A section encompassing the bulbar medulla (M) and cortex (C). Note that ascending blood vessels bound large polygonal areas whose apices anastomose fist and, within the cortex, bound smaller polygonal areas of the neuropil. **b.** High magnification view of the bulbar cortex showing the distribution and polygonal arrangement of anastomotic blood vessels. To note is the progressively larger caliber of blood vessels upwards. **c.** Camera lucida drawing showing the convergence of capillary blood vessels from the rostral migratory stream (RMS) to the glomerular layer (GL). To note is that blood vessels resolve in venous blood vessels within the glomeruli (shaded) by two routes. Namely, by perforating vessels from the RMS (gray arrows) to glomeruli or indirectly, via anastomotic blood vessels. Black arrows = Glomerular veins; arrowheads = venous glomerular sinuses. **d.** Cartoon illustrating a perforating, direct (white) and indirect (black) blood vessel convergence from the bulbar medulla (dark gray) and deep cortex (light gray) to the glomerular layer (GL).

**5.** High magnification view of twin fibers originating fibro-vascular complexes. The video shows first the superficial plane and proceeds throughout a blood vessel outlined in online resource (Fig. 1). To note is that axons and tributary fibro-vesicular complexes are embedded in the astrocytic end-feet that appear discontinuous but homogenously impregnated (light-brown) around the blood vessel’s wall.

**Figure S1.**
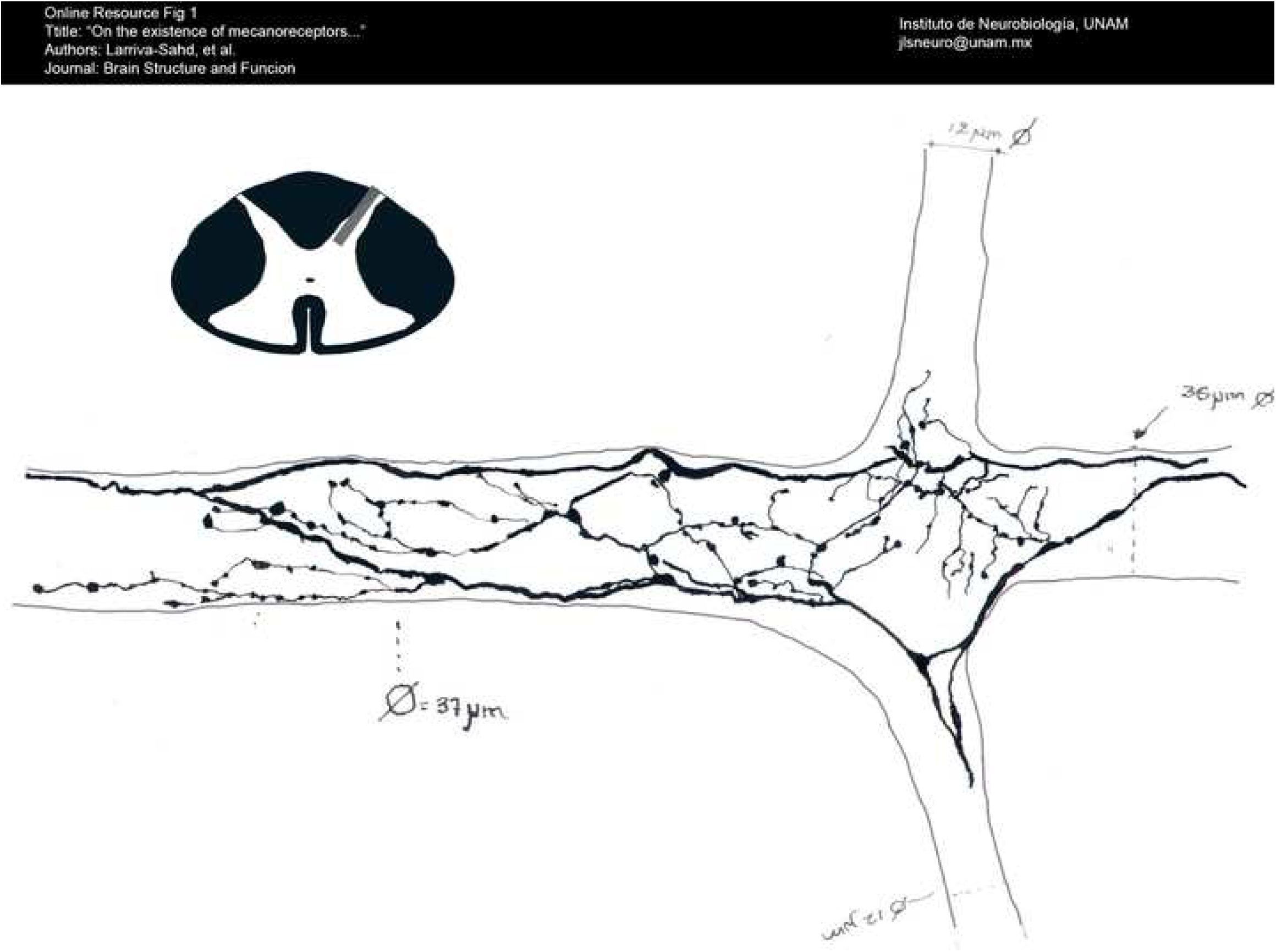

**Figure S2.**
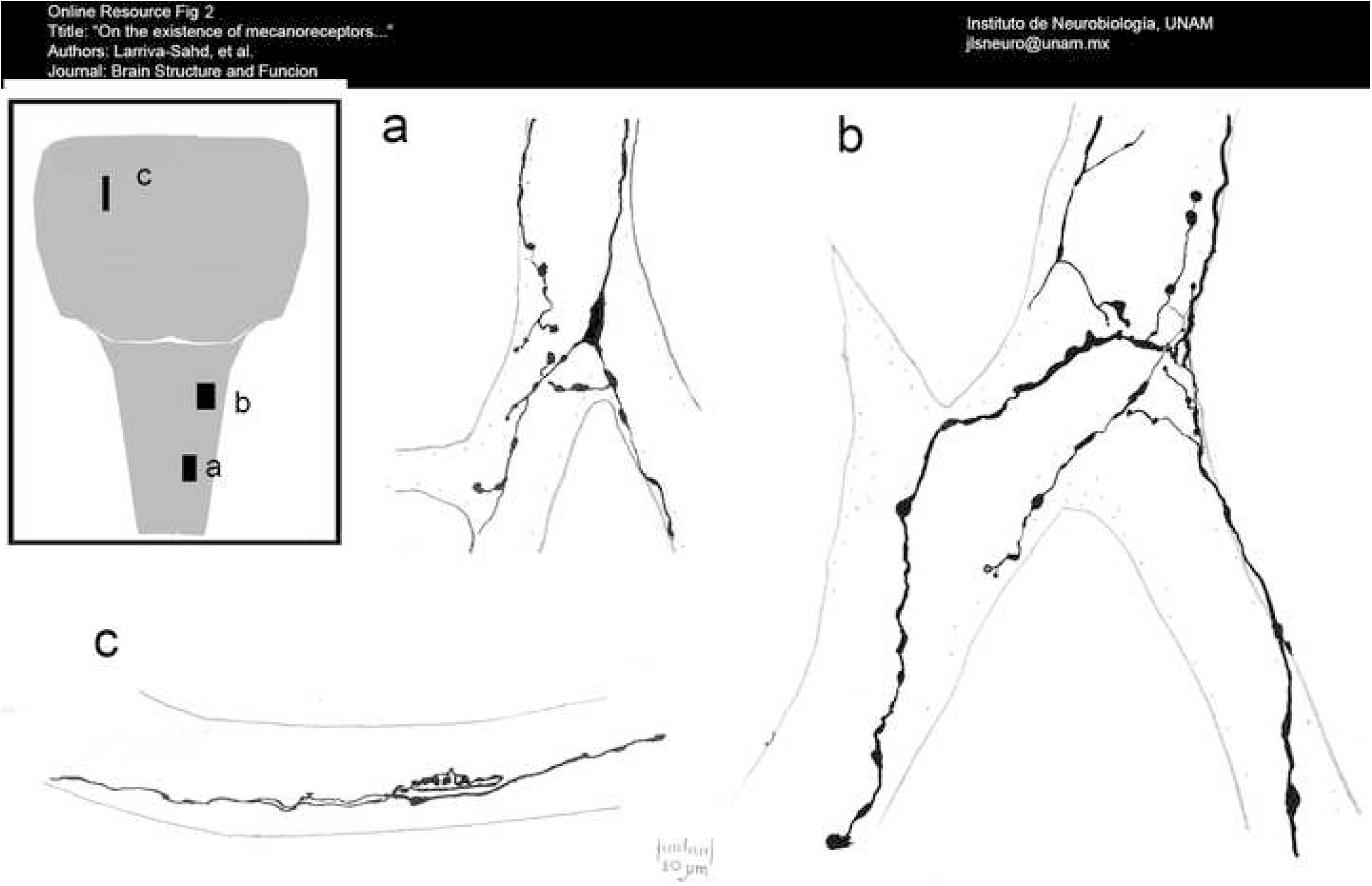

**Figure S3.**
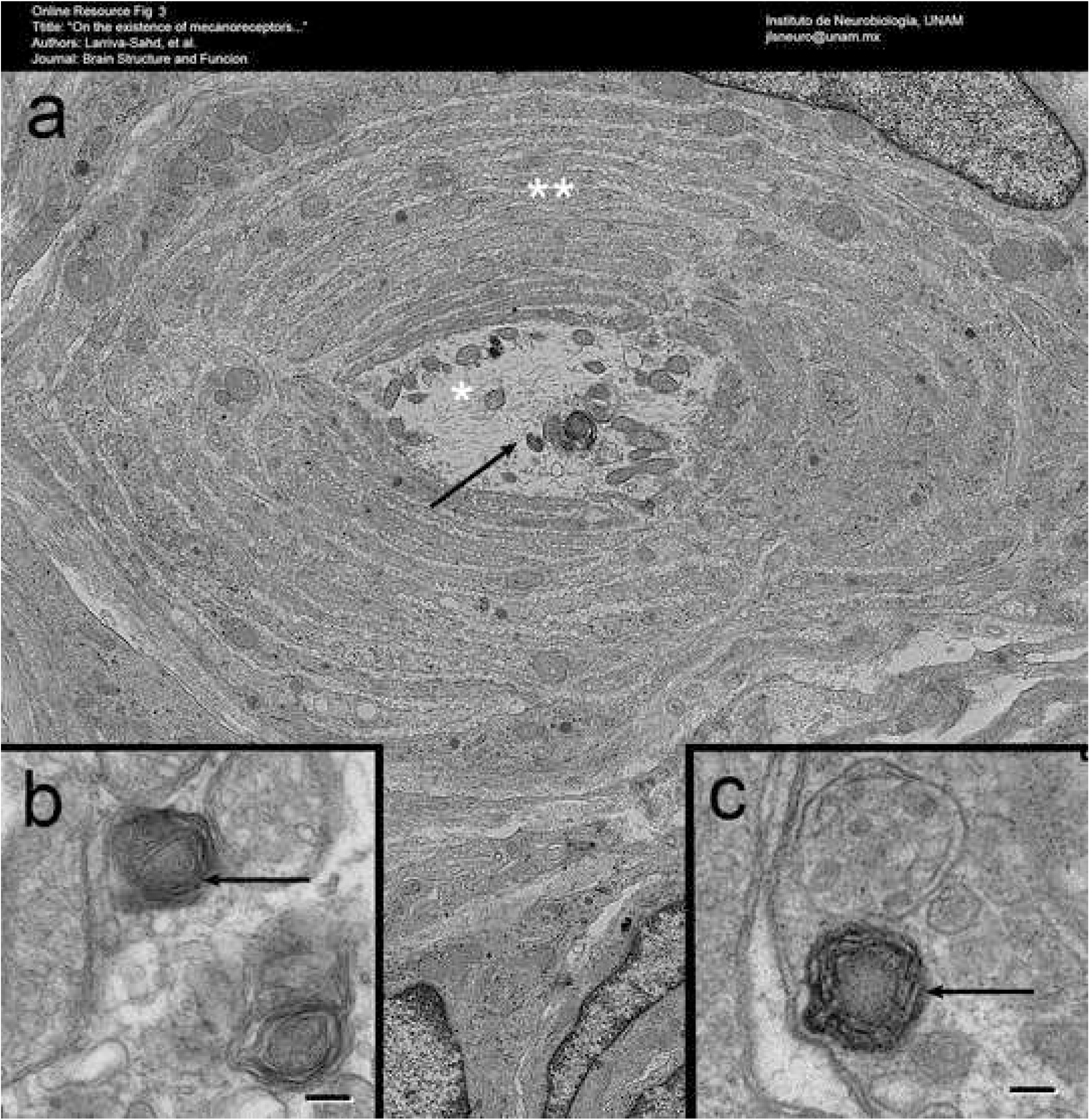

**Figure S4.**
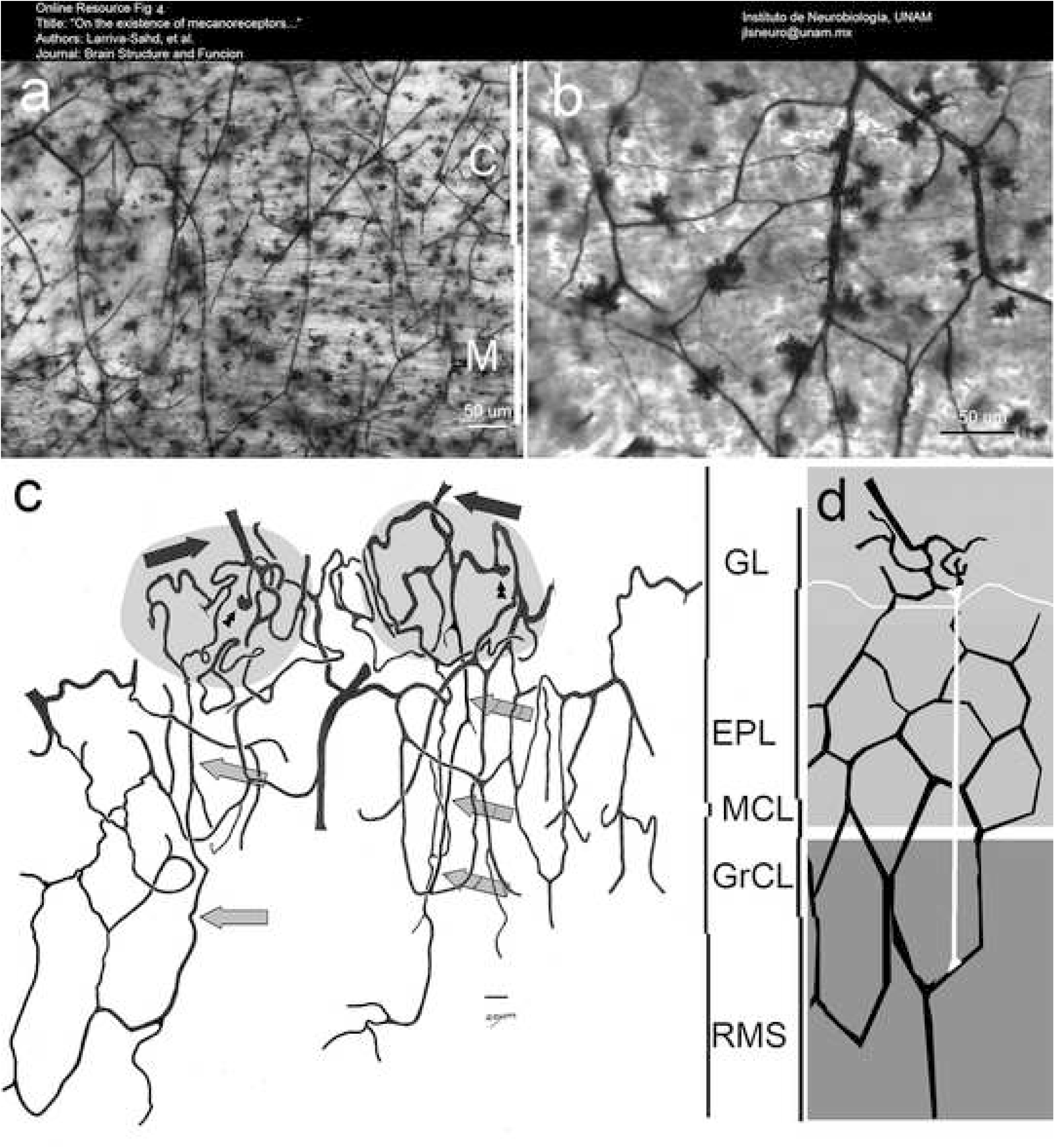

